# Reading Increases the Compositionality of Visual Word Representations

**DOI:** 10.1101/653956

**Authors:** Aakash Agrawal, K.V.S. Hari, S. P. Arun

## Abstract

Reading causes widespread changes in the brain but its effect on visual word representations is unknown. Reading may facilitate visual processing by forming specialized detectors for longer strings, or by making word responses more predictable from single letters, that is by increasing compositionality. We provide evidence for the latter hypothesis by comparing readers and nonreaders of two Indian languages, Telugu and Malayalam. Readers showed decreased interactions between letters during visual search, which predicted their overall reading fluency. Brain imaging revealed increased compositionality in readers, whereby responses to bigrams were more predictable from single letters. This effect was specific to the lateral occipital region, where activations best matched behavior. Thus, reading facilitates visual processing by increasing the compositionality of visual word representations.

## INTRODUCTION

Reading a word involves processing its visual form, associating it with spoken sounds, and processing its overall meaning. Consequently, learning to read alters a variety of brain systems including the visual, auditory and language regions (Dehaene *et al*., 2015). In particular, reading has a profound influence on the visual regions. It leads to the formation of the visual word form area (VWFA) in the fusiform gyrus, which is selectively activated by words of familiar scripts, and by intact words over scrambled controls, and whose activation levels predict reading fluency (Dehaene *et al*., 2015). But reading also causes widespread changes throughout the visual cortex, as shown by the greater activation for intact words relative to scrambled controls (Dehaene *et al*., 2010; Dehaene and Cohen, 2011; Szwed *et al*., 2011) as well as for familiar over unfamiliar scripts (Baker *et al*., 2007; Bai *et al*., 2011; Szwed *et al*., 2014; Krafnick *et al*., 2016).

Despite these insights, several fundamental questions remain unanswered regarding how reading affects letter and word representations. Does reading alter single letter representations? Does it alter word representations over and above the effect on single letters? These questions have been difficult to answer for two reasons: First, letter representations with and without reading expertise are difficult to characterize because many Western languages use the same script, making it difficult to find subjects fluent in distinct scripts without other confounding factors like phonological mapping, writing systems and literacy (Dehaene *et al*., 2015). Indian languages offer a unique opportunity to investigate these issues because of their diverse alphabetic scripts with shared phonological mapping and writing systems (Nag, 2017). This makes it possible to compare subjects proficient in reading distinct scripts while holding constant other confounding factors.

Second, to characterize changes in word representations, it is critical to establish a quantitative model to relate word responses to letter responses. According to an influential account, reading facilitates visual processing through the formation of specialized local combination detectors (LCDs) that detect feature combinations across letters (Dehaene *et al*., 2005). Evidence in favor of this account comes from the increased activation of VWFA as letter strings become orthographically similar to real words (Binder *et al*., 2006; Vinckier *et al*., 2007). However, these results are based on comparing letter strings equated for mean letter frequency. These matched letter strings may contain letters of disparate frequencies or medium frequencies at different positions, which could elicit different responses simply because of letter frequency and position effects (Scaltritti, Dufau and Grainger, 2018). Thus, evidence for LCDs is inconclusive in the absence of a systematic relationship between word and letter responses.

## RESULTS

We compared letter and word representations in distinct groups of readers with similar educational levels but fluent in either of two Indian languages (Telugu and Malayalam). These languages have distinctive scripts with many shared phonemes, and highly similar writing systems (Fig. 1). We selected visually distinct letters with identical pronunciation from both languages (Section S1). This design eliminates not only confounding factors due to phonology, writing systems or literacy, but also isolates the effect of reading expertise from intrinsic shape differences across the two scripts.

**Fig. 1.**
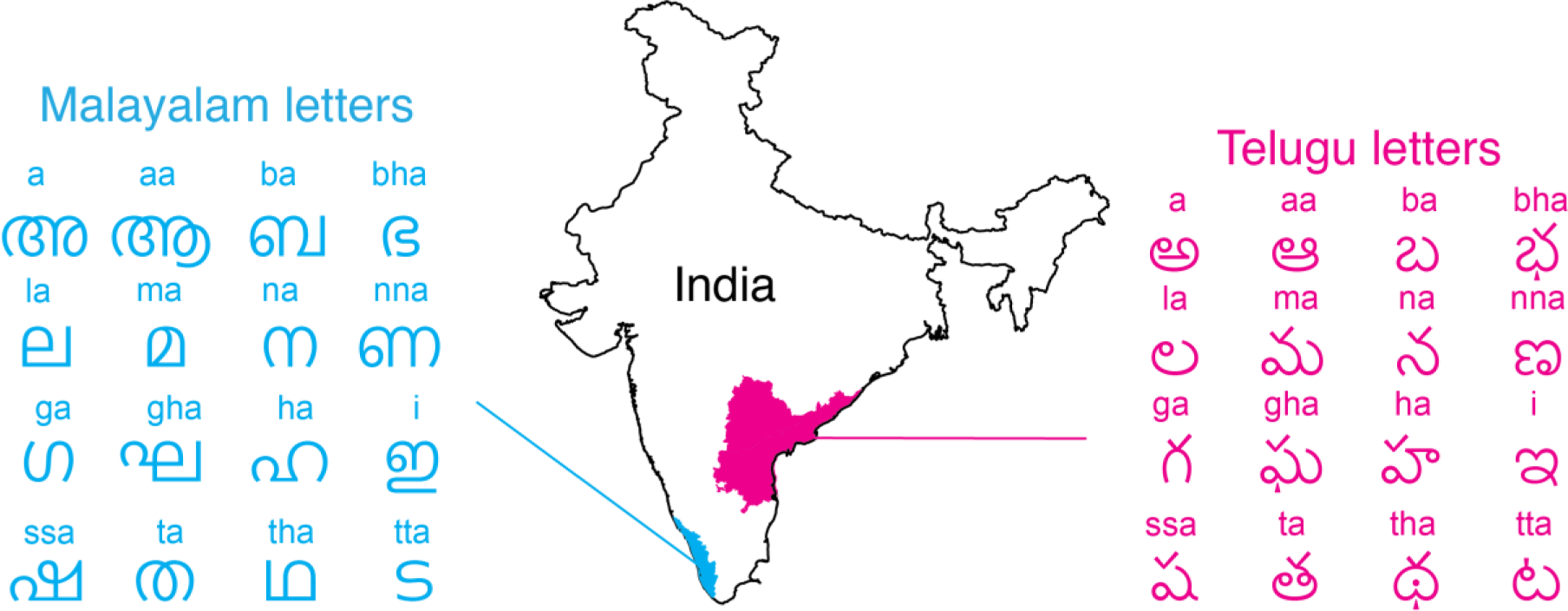
Telugu and Malayalam scripts. The Telugu and Malayalam languages are spoken in geographically distinct regions in India marked in *magenta* and *cyan* respectively (Map courtesy: Free Vector Maps). They have distinct letter shapes but share many phonemes (indicated above each letter). Only 16 example letters are shown here from each language; Telugu has 60 letters and Malayalam has 53 letters. The full set of stimuli are shown in Section S1.

### Single letter searches

We first investigated whether reading expertise modulates single letter representations. We recruited 39 readers (19 Telugu, 20 Malayalam) to perform an oddball visual search task involving Telugu letters and Malayalam letters (Fig. S1). An example search using Telugu letters is shown in Fig. 2A. Subjects were equally accurate on searches involving known and unknown scripts (mean accuracy: 99% for known, 98% for unknown scripts). However, they were faster for searches involving letters of known scripts (Fig. 2B). To compare letter representations, we used the reciprocal of search time as a measure of dissimilarity between letters (Arun, 2012). It can be interpreted as the underlying salience signal that accumulates during visual search (Sunder and Arun, 2016). It combines linearly across object attributes (Pramod and Arun, 2014, 2016), search types (Vighneshvel and Arun, 2013) and even across top-down influences (Sunder and Arun, 2016).

**Fig. 2.**
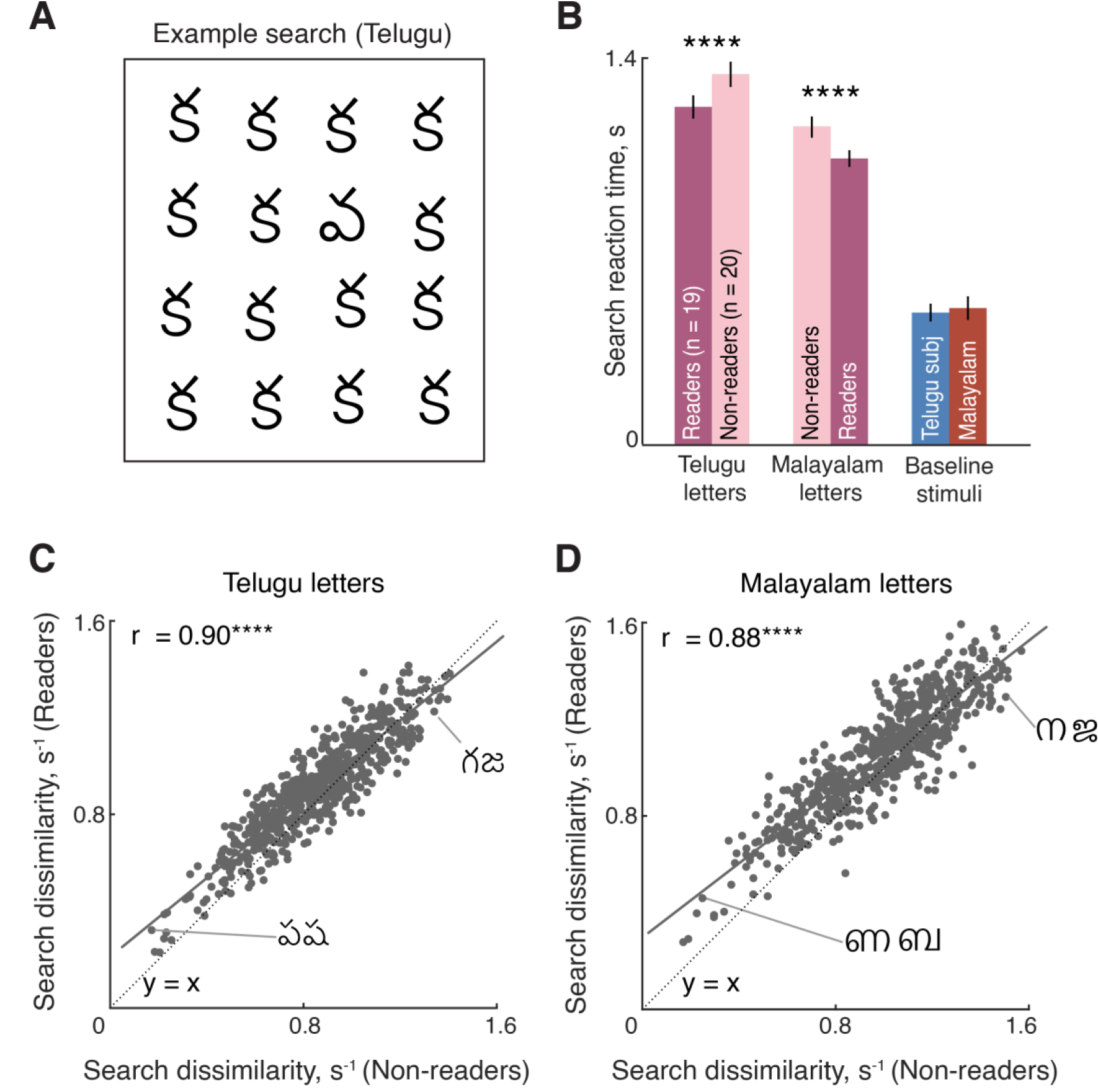
Readers discriminate single letters better. A. Example single letter search array from Experiment 1 using Telugu letters. B. Average search time for readers (*dark*) & nonreaders (*light*) of Telugu and Malayalam letters. The baseline response time is also shown for each subject group. Asterisks indicate statistical significance (**** is p < 0.00005). C. Pairwise search dissimilarity for 630 pairs of Telugu letters plotted for readers against that of non-readers. Each point represents one search pair. An example easy and hard search pair are shown. The dotted line is the y = x line, and the solid line is the best-fitting line to the data. Asterisks indicate statistical significance (**** is p < 0.00005). D. Same as in (C) but for Malayalam letters.

For each language, we plotted the pairwise dissimilarity for readers against that of non-readers across all letter pairs. This revealed a strong positive correlation for both Telugu letters (Fig. 2C) and Malayalam letters (Fig. 2D). These correlations were close to the consistency of the responses within each group (correlation between dissimilarities in odd- and even-numbered subjects: r = 0.83 & 0.87 for readers and non-readers of Telugu; r = 0.83 & 0.87 respectively for Malayalam; all correlations p < 0.00005). Reading expertise also resulted in increased dissimilarity for more similar letters, as shown by a negative correlation between baseline letter dissimilarity (as measured in non-readers) and the increase in dissimilarity for readers over non-readers (r = −0.43 for Telugu & r = −0.49 for Malayalam, both correlations p < 0.00005). These subtle alterations did not affect the global arrangement of letters in perceptual space (Section S2). Letters that co-occurred in a bigram showed greater similarity in readers (Section S2), and their sounds were perceived as more similar (Section S3). In sum, reading subtly alters letter representations through increased discrimination of similar letters.

### Bigram searches

Next we set out to characterize how reading expertise affects the representations of longer strings. Subjects performed oddball visual search involving bigrams of either familiar or unfamiliar scripts (Fig. 3A). Readers were again faster to discriminate bigrams of their known script over unknown scripts (Fig. 3B). These changes had a subtle effect on bigram representations, as shown by a strong correlation between bigram dissimilarities between readers and non-readers (r = 0.80 for Telugu, r = 0.83 for Malayalam, p < 0.00005). These subtle alterations did not result in qualitative changes in the overall perceptual representation (Section S4). Importantly, are these changes driven solely by the increased discrimination of single letters? Or are there additional emergent properties that make readers better able to distinguish bigrams?

**Fig. 3.**
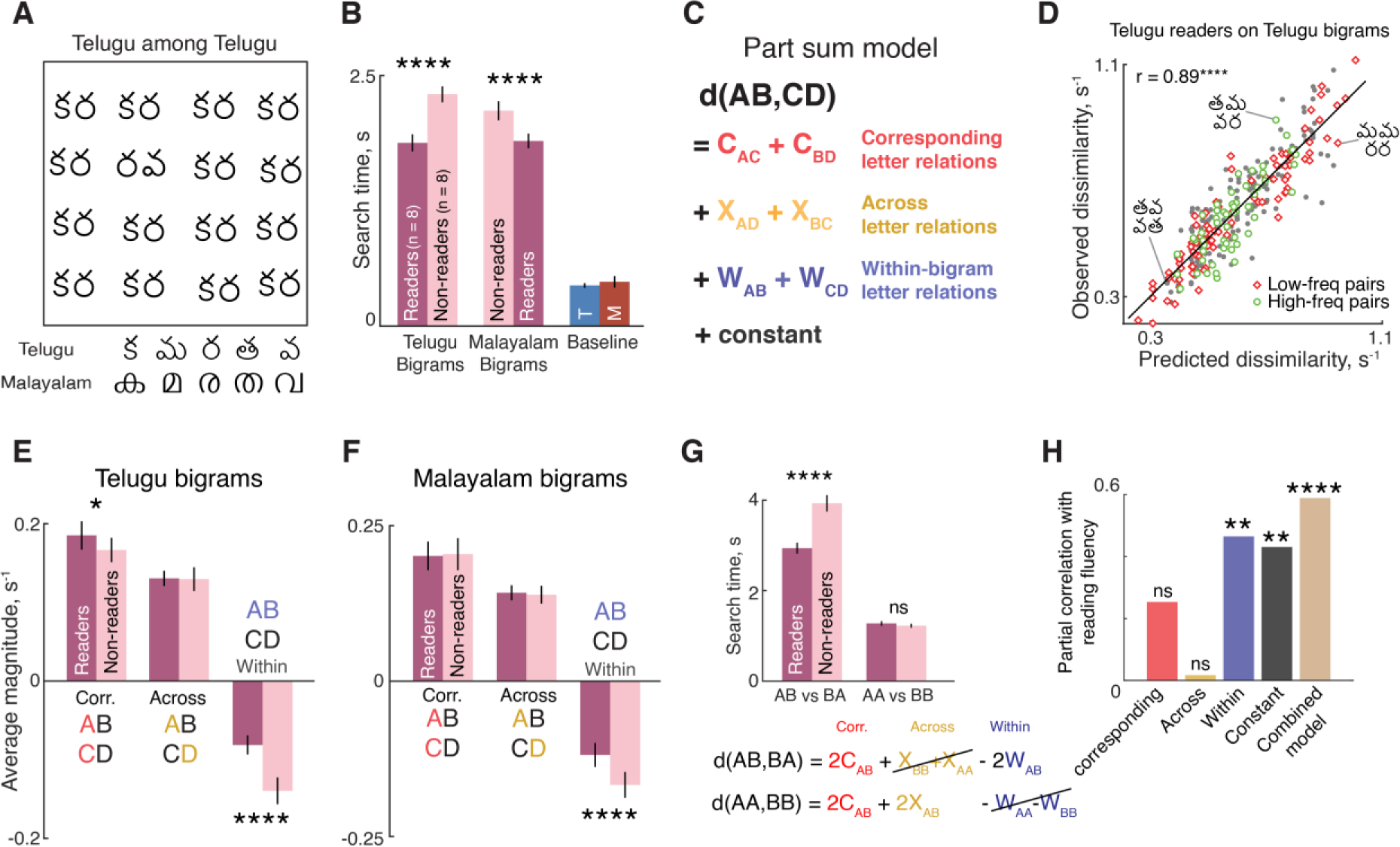
Compositionality for known scripts in readers. A. Example search array using Telugu bigrams (Experiment 2). B. Average search time for readers and non-readers of Telugu bigrams and Malayalam bigrams. Baseline response times are also shown for the two subject groups. Asterisks indicate statistical significance (**** is p < 0.00005). C. Schematic of the Part-sum model. According to this model, the net dissimilarity (1/RT) between bigrams ‘AB’ and ‘CD’ can be explained using single-letter dissimilarities between letters at corresponding locations, opposite locations in the two bigrams, and within each of the bigrams (see text). D. Observed bigram dissimilarity plotted against predicted dissimilarities from the part sum model for Telugu readers on Telugu bigrams. Each point represents one search pair. Searches with infrequent bigrams (n = 91, *red diamonds*) and frequent bigrams (n = 55, *green circles*) are plotted separately from the rest (n = 154, *gray circles*). The dotted line is the y = x line. E. Part sum model parameters (averaged across ^5^C_2_ = 10 part relations) for letter dissimilarities at corresponding locations, opposite locations and within bigrams for readers (*dark*) and non-readers (*light*) for Telugu bigrams. Error bars indicate standard deviation. Asterisks indicate statistical significance (* is p < 0.05, **** is p < 0.00005 on a sign-rank test across 10 part relations between readers and non-readers). F. Same plot as in (E) but for readers and non-readers of Malayalam bigrams. G. Average search reaction time for transposed letter searches (e.g. AB among BA) and repeated letter searches (e.g. AA vs BB) for readers and non-readers (averaged across Telugu & Malayalam readers). Asterisks indicate significant difference (**** is p < 0.00005 on a rank-sum test across search times for 20 AB-BA pairs across the two languages, or across AA-BB pairs). *Top*: Model equations showing how smaller within-bigram terms lead to increased dissimilarity for transposed letters but not repeated letters. For transposed letter searches, letters are identical at opposite locations so the opposite-location terms are multiplied by zero, but the smaller within-bigram terms for readers lead to larger dissimilarities (and therefore faster searches). For repeated letter searches, the within-bigram terms are multiplied by zero by definition and therefore there is no benefit for readers. H. Partial correlation between reading fluency and each part-sum model term (after factoring out all other terms). The combined model is based on predicting reading fluency as a linear combination of all model terms.

To address these issues, we drew upon our finding that dissimilarities between object parts combine linearly in visual search (Pramod and Arun, 2016). Specifically, the net dissimilarity between two bigrams AB & CD is given as a linear sum of part relations at corresponding locations, part relations at opposite locations and part relations within each bigram (Fig. 3C; see Methods). Given many pairwise dissimilarities between bigrams, this part sum model attempts to recover the underlying letter-letter relations that accurately predict this data. The model works because the same letter pair, say AC, is present in many other bigrams at corresponding locations (e.g. AB-CD, AD-CE, BA-DC etc), allowing us to recover its contribution to the overall dissimilarity whenever A and C are present at matched locations in two bigrams. Likewise, the pair AC is present in many bigrams at opposite locations (e.g. AB-DC, AD-EC, etc) and within bigrams (e.g. AC-BD, AC-DE etc) which allows us to recover its contribution to the net dissimilarity when it occurs at opposite locations in two bigrams, or likewise within bigrams. This model (based on 1/RT or search dissimilarity) outperformed other models with fewer parameters, as well as models based on reaction time (Section S4).

This model yielded excellent predictions of the data. It yielded a significant positive correlation between the observed and predicted bigram dissimilarities for Telugu readers tested on bigrams of their script (Fig. 3D). Because model coefficients represent dissimilarities between single letters, we first asked whether they are consistent with each other. This was indeed the case: we found a significant correlation between corresponding terms and across terms (r = 0.81, p < 0.005) and a negative correlation between corresponding and within-terms that approached significance (r = −0.62, p = 0.06). The negative sign of the within-terms represents an effect akin to distracter heterogeneity in visual search (Vighneshvel and Arun, 2013; Pramod and Arun, 2016): when the letters in a target bigram are similar to each other, search for that bigram among distractors is more efficient. All three types of terms contributed to the overall model fit (Section S4). The corresponding terms were correlated with the single letter dissimilarities observed in Experiment 1 (r = 0.83, p < 0.005).

If reading expertise led to the formation of specialized detectors for letter combinations, the part sum model should be unable to predict searches involving high-frequency bigrams, because it only encodes single letter dissimilarities but not bigram frequency. Note that the model can account for letter frequency effects because it estimates the underlying single letter dissimilarity which in turn could depend on letter frequency. We observed no qualitative difference between model fits for high-frequency bigram pairs compared to low-frequency pairs (Fig. 3D). A statistical comparison of the residual error between low- and high-frequency pairs revealed no significant difference (average model residual error: 0.07 for 91 low-frequency pairs, 0.08 for 55 high-frequency pairs; p = 0.96, rank-sum test). We observed similar patterns for readers of Malayalam letters (model correlation = 0.91, p < 0.0005; average residual error: 0.08 for 45 low-frequency pairs; 0.07 for 105 high-frequency pairs, p = 0.06).

The part sum model yielded excellent fits to the observed bigram dissimilarities for both readers and non-readers (model correlations for readers and nonreaders: r = 0.89 & 0.90 for Telugu; 0.91 & 0.92 for Malayalam, p< 0.00005). If model predictions are equally good for readers and non-readers, then what makes readers faster than non-readers? We compared the strength of corresponding, across and within model coefficients for readers and non-readers for Telugu bigrams and Malayalam bigrams. Model coefficients for corresponding and across locations were both positive, which means that dissimilar letters at these locations in the two bigrams lead to larger net 0.89 dissimilarity. However, the larger magnitude of the corresponding terms means that dissimilar letters at these locations have a larger effect. For both languages, corresponding and across model coefficients were similar in magnitude but within-bigram terms were systematically smaller in magnitude for readers compared to non-readers (Fig. 3F-G).

This reduced magnitude for readers results in larger dissimilarities and consequently, easy searches. To confirm that this was indeed the case, we calculated the correlation between the observed difference in reaction times between readers and nonreaders, and asked whether this could be explained by the difference in the respective part sum model predictions for each group. This revealed a positive and statistically significant correlation (r = 0.59 for Telugu bigrams, r = 0.55 for Malayalam bigrams, p < 0.00005).

We also confirmed that the first letter in the bigram is more salient than the second, consistent with the first letter advantage observed in letter recognition tasks (Section S4).

If reading does indeed reduce letter interactions within a bigram, then it should have no effect on bigrams with identical letters, since the within-bigram dissimilarity is zero by definition. Therefore we predicted that the dissimilarity between repeated letter bigrams such as AA & BB should not be different for readers and non-readers. In contrast, the dissimilarity between the transposed bigrams AB and BA should be strongly influenced by reading expertise because the within-bigram terms are non-zero while the across-location terms are zero. Thus, the part-sum model predicts that readers should be faster than nonreaders on transposed letter searches (AB-BA) but not repeated letter searches (AA-BB), even though these both types of searches differ in two letters.

This was indeed the case: Readers were faster on transposed bigram searches over non-readers (Fig. 3G). However, they were equally fast for repeated letter searches (average search times for readers and non-readers: 1.29 & 1.30 s for Telugu letters, p = 0.91; 1.23 & 1.12 s for Malayalam letters, p = 0.62; Fig. 3G). The lack of effect for repeated letter searches was not due to a floor effect, because there were many easier searches for both readers and non-readers (least average search time for readers and non-readers: 0.90 & 0.89 s for Telugu letters, 0.79 & 0.80 s for Malayalam). Thus, reading expertise produced increased discrimination of letter transpositions compared to repeated letters, and this effect is due to decreased letter-letter interactions within a bigram. We also tested subjects on visual search for trigrams in a separate experiment. Here too, the part sum model yielded excellent fits, with reduced within-trigram letter interactions for readers compared to non-readers (Section S5).

### Bigram dissimilarity and reading fluency

If within-bigram letter interactions are smaller for readers over non-readers, could these interactions predict reading fluency? To investigate this we estimated model parameters using the pairwise bigram dissimilarity from each subject (see Methods), and asked if this predicted reading fluency. To be sure that the contribution of each term was independent of the others, we performed a partial correlation analysis. This revealed a significant partial correlation only for within-bigram interactions and the constant term but not the others (Fig. 3H; Section S6). In other words, subjects with weaker within-bigram interactions were faster at reading. Likewise, subjects with faster motor responses (i.e. larger constant term) were also faster at reading. Combining these factors together yielded a better model fit than each separately, suggesting that they exert distinct influences on reading (Fig. 3H; Section S6).

### Brain imaging of single letters and bigrams

We have shown that reading subtly altered letter representations by making similar letters more discriminable and by reducing interactions between letters within a bigram. However, these results were based on simultaneously shown pairs of bigrams and the fact that interactions decrease within a bigram was only an indirect inference. In Experiment 3, we sought to directly quantify whether letters interact in a bigram by presenting one stimulus at a time and by relating bigram responses to single-letter responses. If reading decreases letter interactions, the responses to bigrams should be more predictable from single letters in readers compared to non-readers. By contrast, if reading leads to specialized bigram detectors, the responses to bigrams should be less predictable from single letters in readers.

A total of 35 subjects (17 Telugu & 18 Malayalam) participated in this experiment. We defined a number of regions of interest (ROI) for further analysis: early visual cortex (V1-V3), V4, Visual Word Form Area (VWFA), Temporal Gyrus (TG) and the Lateral Occipital region (LO). All ROIs were defined using a combination of anatomical considerations and functional localizers (see Methods). A representative subject brain with these ROIs is shown in Fig. 4A. We also performed equivalent searchlight analyses (Section S7). In the main experiment, subjects viewed single letters and bigrams while performing a 1-back task.

**Fig. 4.**
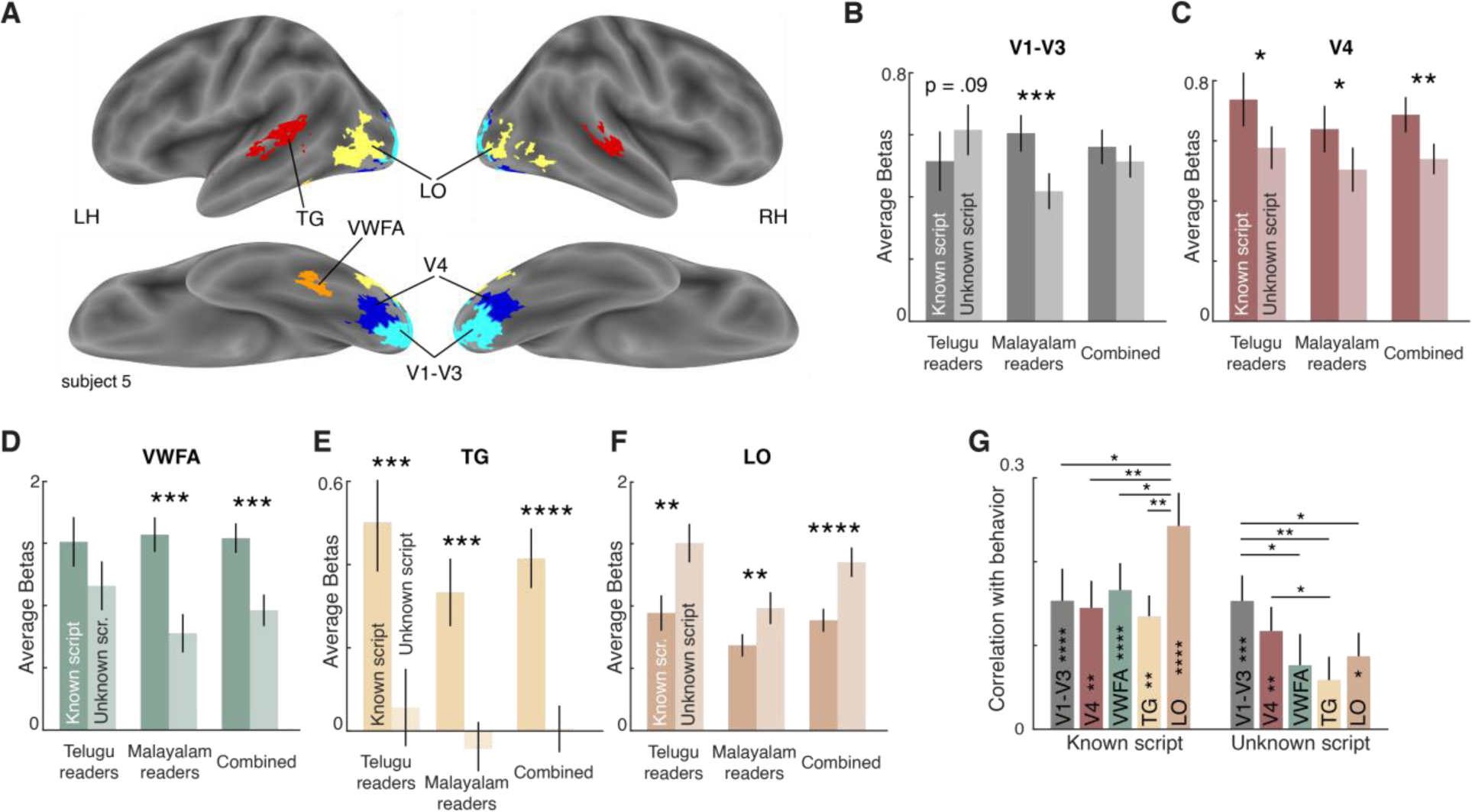
Neural correlates of reading expertise. A. ROIs for an example subject, showing V1–V3 (*cyan*), V4 (*blue*), LO (*yellow*), VWFA (*orange*) and TG (*red*). B. Average activation levels for known and unknown scripts for Telugu readers, Malayalam readers and combined data for V1-V3. Error bar indicate ±1 SEM across subjects. Asterisks indicate statistical significance (* is p < 0.05, ** is p < 0.005, etc. in a sign-rank test comparing subject-wise average activations). C-F. Same as in B but for V4, VWFA, TG and LO respectively. G. Correlation between neural dissimilarity in each ROI with behavioral dissimilarities for bigrams of the known script (*left*) and unknown script (*right*). Error bars indicate standard deviation of the correlation between the group behavioural dissimilarity and ROI dissimilarities calculated repeatedly by resampling of subjects with replacement across 1000 iterations. Asterisks along the length of each bar indicate statistical significance of the correlation between group behaviour and group ROI dissimilarity (* is p < 0.05, ** is p < 0.005, etc). Asterisks spanning bars indicate the fraction of bootstrap samples in which the observed difference was violated (* is p < 0.05, ** is p < 0.005 etc). All significant comparisons are indicated.

### Do known and unknown scripts elicit differential activations?

We first compared overall activation levels in each ROI. For each subject, we calculated the average activation across all voxels and across all stimuli within each script (known and unknown). We compared subject-wise activation levels between known and unknown scripts (Fig. 4B-F).

For early visual areas (V1-V3), we observed opposite effects for known and unknown scripts for Telugu and Malayalam readers, suggesting that Malayalam letters activate early visual areas more than Telugu letters. This comparison was statistically significant (average activations of V1-V3: 0.61 for Telugu, 0.47 for Malayalam, p < 0.00005 using a Wilcoxon signed rank test on subject-wise activations). This difference was highly significant in Malayalam readers (p < 0.0005) but approached statistical significance in Telugu readers as well (p = 0.09). We surmised that Malayalam letters activate early visual cortex better because they are generally wider compared to Telugu letters. We therefore compared both the overall width and ink area (number of pixels) between Telugu and Malayalam letters. As we expected, Malayalam letters were significantly wider and had significantly more ink per letter compared to Telugu letters (mean ± std of width in pixels: 160 ± 37 for Telugu and 191 ± 34 for Malayalam, p = 0.096, rank-sum test; average fraction of non-zero pixels: 0.08 for Telugu & 0.11 for Malayalam; p < 0.005, rank-sum test across letters used in this experiment). The activation of early visual cortex is thus driven by low-level differences in letter shape.

We obtained identical trends in V4, VWFA and TG: known scripts consistently elicited greater activations in readers of both languages (Fig. 4C-E). A searchlight analysis confirmed these trends, but additionally revealed that this trend along a nearly continuous swath of cortex along the ventral surface from V4 to the VWFA, as well as in several regions around the temporal gyrus (Section S7). We observed an opposite pattern in the LO. Here, known scripts elicited weaker activation compared to unknown scripts for both languages (Fig. 4F). A searchlight analysis revealed that this was true on the dorsal portion of the occipitotemporal cortex as well as in parietal regions (Section S7). Reading expertise thus leads to widespread changes specifically in high-level visual areas, but with opposite effects in LO compared to V4/VWFA.

We performed two additional analyses using overall activation levels. First, there were differences in overall activation with bigram frequency for Telugu but not Malayalam, but this effect was abolished upon factoring out letter frequency effects (Section S7). Second, we observed a positive correlation between mean VWFA activation levels and reading fluency across subjects (Section S7), concordant with other studies (Dehaene *et al*., 2015).

### Neural correlates of behavior

Having obtained systematic effects of reading on single letter and bigram dissimilarities in Experiments 1-2, we looked for the neural correlates of these effects by calculating neural dissimilarities for each ROI. Specifically, for each ROI in a given subject, we calculated the neural dissimilarity between pairs of images using the correlation distance between the voxel activations of the two images (1-r), and averaged this dissimilarity across subjects. In this manner, we calculated average pairwise neural dissimilarities for all pairs of stimuli in each ROI (for the dissimilarity matrices, see Section S7). We performed a separate experiment (Experiment 4) to estimate the pairwise dissimilarities for the bigrams used in fMRI. We then asked how well these pairwise dissimilarities matched behavior for known scripts and unknown scripts. The results revealed two interesting patterns. First, neural dissimilarities in a number of areas were significantly correlated with behavior for both known and unknown scripts (Fig. 4G). However, the best match with behavior for known scripts was in LO, whereas for unknown scripts it was in V1-V3. A searchlight analysis confirmed these trends (Section S7): dissimilarities for known bigrams best matched with neural dissimilarities in occipitotemporal cortex centred around LO, but also with the activation of parietal and motor regions. In contrast, the dissimilarities for unknown bigrams best matched the neural dissimilarities in early visual areas. Thus, reading expertise biases visual processing towards higher visual areas.

### Does reading alter the compositionality of bigram representations?

We next turned to the critical question of whether reading alters the compositionality of bigram representations. We devised a model to predict the response of a population of voxels to a given bigram using a linear sum of the population response to each individual letter in the bigram (Fig. 5A). This resulted in a separate population model for each bigram. Doing so allows the model to estimate the average compositionality across a population of correlated voxels and overcome the inherent noise in individual voxels. To evaluate model fits, we compiled model predictions for each voxel across bigrams and compared it with its observed activations. Doing so prevents the model fits from being biased by voxels with large activation levels. We obtained similar results upon fitting a separate model to each voxel. We compared the average model fit for each subject in a given ROI for known and unknown scripts. We obtained comparable model fits for known and unknown scripts in most ROIs (Fig. 5B). The sole exception was LO, where bigrams of known scripts were better predicted by single letter responses compared to bigrams of unknown scripts (Fig. 5B). A searchlight analysis revealed that this effect was localized to the anterior portion of left LO and to the right fusiform gyrus (Section S7). Since these regions were identified using their higher model fit for known scripts, any direct comparison of model performance would constitute double-dipping. To avoid this circularity, we performed a split-half analysis. We identified the anterior portion of LO using odd-numbered subjects, and compared the model fits in even-numbered subjects, and vice-versa. This revealed significantly larger model fits in anterior LO for known compared to unknown scripts for Telugu and Malayalam separately as well as in both languages combined (Figure 5C). We obtained similar results in the right fusiform gyrus (Section S7).

**Fig. 5.**
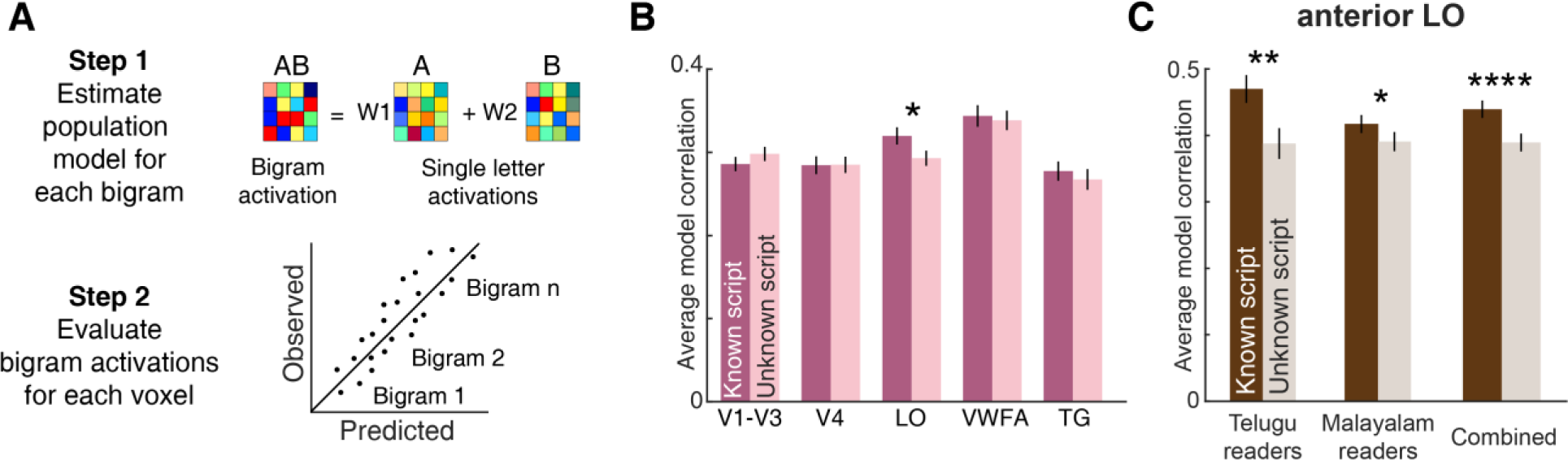
Compositionality of neural bigram representations. A. Schematic of the voxel population model. The response of each bigram across voxels is modelled as a linear combination of the constituent letter responses. To evaluate the model fit, we calculated the correlation between observed and predicted activations for each voxel. B. Average model correlation (across voxels) for each ROI for known (*dark*) and unknown (*light*) scripts. Error bar indicate SEM across subjects. Asterisks indicate statistical significance (* is p < 0.05 using a sign-rank test on subject-wise model correlations between the two groups). C. Average model correlation in the anterior LO for known (*dark*) and unknown (*light*) scripts for Telugu and Malayalam languages separately as well as both languages combined. Asterisks represent statistical significance, as obtained using a sign-rank test comparing average model correlations across subjects (* is p < 0.05, ** is p < 0.005 and so on).

The increased compositionality for bigrams in object-selective cortex could be an incidental artefact of having stronger signal levels overall, which could increase the explainable variance and therefore model performance. However, this is unlikely because known scripts evoke weaker activity in LO, which should have led to weaker, not stronger model predictions.

We conclude that reading increases the compositionality of bigram representations specifically in object-selective cortex.

## DISCUSSION

We investigated the effect of reading expertise on letter representations by comparing Telugu and Malayalam readers using a combination of behavior and brain imaging. In behavior, subjects discriminated letters of their known script better compared to unknown scripts. This is consistent with the increased discrimination of familiar targets observed for natural objects in visual search (Mruczek and Sheinberg, 2005). We found that the net dissimilarity between strings (bigrams/trigrams) can be accurately predicted using pairwise dissimilarities between letters in the two strings. This is consistent with our previous study where we reported this for objects (Pramod and Arun, 2016). This model was able to predict virtually all the explainable variation in the search data for both readers and non-readers. Importantly, these changes in visual processing directly predicted reading fluency in readers.

If reading expertise led to the formation of specialized bigram or trigram detectors, our models, based only on single letter responses, would have shown worse performance for known scripts than for unknown scripts, and for frequent over infrequent bigrams. We found no such effects. Thus, bigram detectors, even if present, do not contribute substantively to the observed effects. Importantly, we were able to precisely quantify the effect of reading on word representations by analysing how model parameters varied between readers and non-readers. Our main finding is that reading expertise made single letters more discriminable, and reduced interactions between letters in a string. Our model accounts for both letter similarity and letter interactions, thereby providing a framework to compare effects of letter substitution and transposition, both widely used as measures of orthographic processing (Grainger *et al*., 2012; Ziegler *et al*., 2013; Dehaene *et al*., 2015). Further, the reduced interactions may work at multiple scales: for instance parsing conjoined frequent words is easier than parsing conjoined infrequent words (e.g. readingdifficulty is easier than heliumchromate). We propose that visual search using letter strings can be a natural, effective and objective measure to study how reading alters visual representations.

Our brain imaging experiment further elucidated the neural basis of word representations. Our main finding is that the anterior ventral portion of LO is a likely locus for the effects observed in behavior. This is because (1) the neural representation of bigrams in LO matched best with behavior for readers; and (2) bigram responses were better predicted from single letters for known scripts specifically in the LO but not in other regions. The former finding is interesting because it suggests that reading shifts the neural basis of behaviour from lower to higher visual areas. The latter finding is interesting because it indicates that reading makes visual processing more efficient by making words easier to parse into letters. We speculate that this may enable parallel sequencing of speech motor movements.

That LO could play a role in reading is consistent with evidence that alexia also induces general visual processing deficits (Behrmann, Nelson and Sekuler, 1998; Roberts, Lambon Ralph and Woollams, 2010; Starrfelt, Habekost and Gerlach, 2010) and often involves damage to regions posterior to VWFA (Barton, 2011; Seghier *et al*., 2012). It is also possible that this effect is present in other areas such as VWFA but undetectable due to having far fewer voxels and therefore weaker statistical power. A conclusive demonstration that anterior ventral LO participates in reading would require perturbing its activity during reading tasks.

We also found a widespread effect of reading expertise across many high-level visual regions, in keeping with the existing literature (Dehaene *et al*., 2010; Szwed *et al*., 2012, 2014). But unlike previous studies, we compared readers of closely related Indian languages with distinct orthographies, shared phonemes and with similar educational levels. This enabled us to establish that these effects were truly due to reading expertise and not due to letter shapes or other confounding factors. We found opposite trends in different areas. In V4 and along the fusiform gyrus up to the VWFA, we found greater activation to known scripts. This is consistent with the effects of learning observed in these regions (Folstein *et al*., 2015; Clarke *et al*., 2016; Skeide *et al*., 2017). In the occipitotemporal regions in and around LO, we observed greater activation for unknown scripts. This is consistent with the increased response to novel stimuli in the homologous region in the monkey, the inferior temporal cortex (Mruczek and Sheinberg, 2007; Meyer *et al*., 2014). These effects could also arise from different effects of attention on these regions, although we note that such attentional effects have never been proposed or reported. Distinguishing familiarity effects from attentional effects is challenging because it required careful independent control of task difficulty, attention and familiarity.

Our observations both confirm and extend our understanding of the VWFA in several ways. First, we consistently localized the VWFA for both Indian languages, and observed no difference in its anatomical location across language (Section S7). This is consistent with other studies where VWFA was observed at similar locations for multiple languages (Bai *et al*., 2011; Szwed *et al*., 2014; Krafnick *et al*., 2016). Second, we found a positive correlation between VWFA activation levels and fluency (Dehaene *et al*., 2015). Third, neural dissimilarity in the VWFA was significantly correlated with behavioral dissimilarity only in readers (Fig. 4G). There have been surprisingly few studies in this regard. Only one study has shown VWFA representations to be correlated with subjective visual dissimilarity (Rothlein and Rapp, 2014), but this could be due to explicit letter reading by subjects. Our measure of behavioral dissimilarity (visual search) did not require explicit reading, and was similar for readers and non-readers. Thus the positive correlation observed in our study implies that VWFA receives letter shape information only for known scripts. Finally, we observed concordant effects in the VWFA and temporal gyrus regions, indicative of its status as an intermediate region between visual and auditory areas (Friederici and Gierhan, 2013; Dehaene *et al*., 2015).

## MATERIALS AND METHODS

All subjects had normal or corrected-to-normal vision and gave written informed consent to experimental protocols approved by Institutional Human Ethics Committee of the Indian Institute of Science. Subjects had similar educational status: they were all undergraduate or graduate students at the Indian Institute of Science. All subjects were also fluent in English, and were fluent in reading either Telugu or Malayalam but not both.

### Fluency test

Subjects were asked to perform a brief fluency test along with every experiment. In this test, a passage of text was shown to the subject in his known script (Telugu or Malayalam). In both languages, this passage described how the head of an Indian village introduced computers to the village and employed software professionals to train residents. This passage was prepared by translating the same English passage into both languages. Subjects were asked to silently read the passage presented on a computer screen and press a button after they finished reading it. After this a dialog box appeared and subjects were asked to summarize the passage in English, and this summary was reviewed offline (by AA) to confirm that the subjects indeed comprehended the passage. The time taken by subjects for the button press was taken as a measure of reading fluency. All but two subjects from Experiment 3 participated in the fluency test. A minority of the subjects (n = 4) declared afterwards that they read the passage multiple times to memorize it, so their data was excluded from subsequent fluency analyses.

### Experiment 1 (Single letters)

A total of 39 subjects (28 males, 25 ± 4 years, 19 Telugu, 20 Malayalam) – participated in this experiment. Here and in all visual search experiments, we chose this sample size because previous studies from our group have obtained highly consistent data using similar sample sizes (Pramod and Arun, 2016). The stimuli consisted of 36 single letters each from Telugu and Malayalam language (shown in Section S1). The font “Nirmala UI” was used as it had uniform stroke width. Subjects performed a baseline motor response task and an oddball visual search task.

In the baseline task, a circle appeared to the left or right of the screen and subjects had to indicate the side on which the circle appeared using a key press (‘Z’ for left, ‘M’ for right). The average response time across 20 trials was taken as a measure of the baseline motor speed (depicted in Fig. 2B). In the visual search task, each trial began with a fixation cross for 500ms followed by 4×4 search array which contained one oddball target and 15 identical distractors (Fig. 2A). The exact position of each item was jittered on each trial according to a uniform distribution with range ±0.25° in the vertical and horizontal directions. This was done to avoid alignment cues from influencing search. The vertical dimension of all letters subtended 2° of visual angle on the screen, and the longer dimension varied depending on the letter. A vertical red line divided the screen into two halves. All stimuli were presented using custom scripts written in MATLAB running the Psychophysics Toolbox (Brainard, 1997). Subjects were instructed to locate the target as fast and as accurately as possible and respond via key press ‘Z’ (for left) and ‘M’ (for right). The trial timed out after 10 s after which the trial was repeated after a random number of other trials. All stimuli were presented in white against black background. In all, subjects completed two search trials corresponding to all ^36^C_2_ pairs of letters in each language, which amounted to 2520 correct trials (^36^C_2_ pairs x 2 languages x 2 repetitions). Incorrect or missed trials appeared randomly later in the task. Only correct responses were analysed. To remove outliers in response times, any response exceeding 5 seconds was removed from analysis provided such a response occurred in less than 15% of the subjects. This step improved data consistency overall, and was done for all behavioral experiments. We obtained qualitatively similar results without this step.

### Experiment 2 (Bigrams)

A total of 16 subjects (10 males, 24 ± 2 years, 8 Telugu, 8 Malayalam) – participated in this experiment. The stimuli consisted of 25 bigrams each from Telugu and Malayalam, created using all possible combinations of 5 single letters (shown in Section S1). The single letters were chosen such that the full stimulus set contained a few frequent bigrams in each language. In all, subjects performed searches corresponding to all possible pairs of the 25 bigrams, which amounted to 1200 correct trials (^25^C_2_ searches x 2 languages x 2 repetitions). All other details were identical to Experiment 1.

### Part-Sum model

Search response times (RT) were averaged across repetitions and subjects to obtain a composite measure which we then converted to a dissimilarity measure (1/RT) as in our previous studies. This resulted in a total of ^25^C_2_ = 300 pairwise dissimilarities between all possible pairs of bigrams. Using the approach reported in our previous study (Pramod and Arun, 2016), we modelled the pairwise dissimilarity between two bigrams AB & CD using as a linear sum of pairwise dissimilarities between single letters at various locations. Specifically:

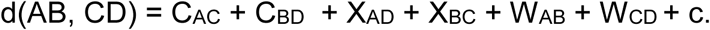

where C_AC_ & C_BD_ represent the contribution distance between letters at corresponding locations in the two bigrams, X_AD_ & X_BC_ represent the distances between letters at opposite locations in the two bigrams, W_AB_ & W_CD_ represent distances between letters within each of the two bigrams and c is a constant term.

This part-sum model is extremely general in that it assumes no systematic relation between single-letter distances at corresponding or across locations in a bigram, or within a given bigram. It works because a given letter pair occurs repeatedly across bigram pairs (e.g. the pair A-C is present at corresponding locations in the bigram pairs AB/CD, AD/CE, EA/DC etc). Because there are 5 unique single letters, there are ^5^C_2_ = 10 single letter distances for each term type (corresponding, across, within), which amounts to a total of 31 parameters (10 of each type x 3 types + 1 constant). Since there are 300 dissimilarity measurements and only 31 parameters, the model parameters can be uniquely estimated from the data. When the above model equation is written down for all 300 pairwise dissimilarities, the set of simultaneous equations can be written down as y = Xb where y is a 300 x 1 vector containing the observed dissimilarities, X is a 300 x 31 matrix with either 0, 1 or 2 as entries depending on the absence, presence or repetition of a particular letter-pair at either corresponding locations, across locations or within the two bigrams of a given pair, and b is a 31 x 1 vector of unknowns. We estimated the model parameters using standard linear regression (*regress* function in MATLAB).

### Experiment 3 (fMRI)

A total of 35 subjects (31 males, 25 ± 3 years, 17 Telugu, 18 Malayalam) participated in functional localizer runs (n = 2) and event-related runs (n = 8) which were randomly interleaved. An anatomical scan was also included for each subject at the beginning. We chose this sample size because it is similar to the number of subjects used in previous studies of reading (Baker *et al*., 2007).

In the functional localiser runs, subjects viewed 16 s long blocks of Telugu, Malayalam, English, scrambled words, objects and scrambled objects while performing a one-back task throughout. In each block, 14 stimuli were randomly selected from a pool of images. The Telugu pool comprised 8 two-letter and 38 three-letter words, and the Malayalam pool comprised 12 two-letter and 38 three-letter words. The English pool comprised 36 four-letter words and 45 five-letter words. Telugu and Malayalam letters were typically wider than English letters, so we used longer English words so that the overall width of the image was roughly equal for all three languages. Each word was divided into grids of dimension 8×4 (for Indian languages) or 8×3 (English) and scrambled words were creating by randomly shuffling the grid. The objects pool comprised 80 man-made objects. Scrambled objects were created by scrambling the phase of the Fourier-transformed images and then reconstructing the phase-scrambled image using the inverse Fourier transform. All images were presented against a black background. Each block consisted of a total of 16 stimuli presented for 0.8 s with a 0.2 s blank interval, among which two randomly chosen stimuli were repeated. Each block ended with a fixation cross presented for 4 s against a blank screen. Thus each block lasted 20 seconds. The size of the object images was about 4.5° along the longer dimension, while the vertical size of the word stimuli was 2.5°, same as in the event-related runs. There were six repetitions of each block across 2 runs and each run lasted for 370s. Stimuli were presented using a custom MATLAB scripts written using Psychophysics Toolbox (Brainard, 1997).

In the event-related runs, the stimuli consisted of 10 single letters and 24 bigrams each in Telugu and Malayalam, amounting to a total of 68 stimuli. The height of the stimuli were equated to subtend 2.5° of visual angle, with longer dimension that varied across stimuli. The bigrams were chosen such that each letter appeared at least 4 times, the mean bigram dissimilarities were similar across the two languages and comprised both high- and low-frequency bigrams (see Section S1 for all stimuli). On each trial, stimulus was presented at the centre of the screen with black background for 300ms followed by a blank screen with fixation cross for 3.7s. In each run, all stimuli were presented once. Subjects were instructed to maintain fixation on the cross and perform a one-back task i.e. to respond using a button press whenever an image repeated twice consecutively. Each run contained eight trials with only fixation cross in order to jitter the inter-stimulus interval and eight randomly chosen images were repeated in a given run. Each run lasted 386 s and there were 8 runs in all, giving eight repeats per stimulus. Post-hoc analyses revealed that subjects were slightly more accurate on performing the task for letters of their known scripts compared to unknown scripts (average accuracy for known and unknown scripts: 88% & 82% for Telugu readers, p < 0.05 using a sign-rank test on accuracy across all subjects; 87% and 67% for Malayalam readers; p < 0.0005). This difference in accuracy presumably reflects facilitated visual processing or increased attention to familiar scripts, but do not by themselves explain the key results from this experiment.

### Data acquisition

Subjects viewed images projected on a screen through a mirror placed above their eyes. Functional MRI data was acquired using 32-channel head coil on a 3T Siemens Skyra at the HealthCare Global, Bengaluru. Functional scans were performed using a T2*-weighted gradient-echo-planar imaging sequence with the following parameters: TR = 2s, TE = 28ms, flip angle = 79°, voxel size = 3×3×3 mm^3^, field of view = 192×192 mm^2^, and 33 axial-oblique slices for whole brain coverage. Anatomical scans were performed using T1-weighted images with the following parameters: TR = 2.30s, TE = 1.99ms, flip angle = 9°, voxel size = 1×1×1 mm^3^, field of view = 256×256×176 mm^3^.

### Data pre-processing

The raw functional MRI data were pre-processed using SPM12 (https://www.fil.ion.ucl.ac.uk/spm/software/spm12/). Raw images were realigned, slice-time corrected, co-registered to the anatomical image, segmented and normalized to the MNI305 anatomical template. Repeating the key analyses with voxel activations estimated from individual subjects yielded qualitatively similar results. Smoothing was performed only on the functional localiser blocks using a Gaussian kernel with FWHM of 5 mm. Default SPM parameters were used and voxel size after normalization were retained to 3×3×3 mm^3^. The data was further processed using GLMdenoise v1.4 (Kay *et al*., 2013). GLMdenoise improves the signal-to-noise ratio in the data by regressing out the noise estimated obtained from task-unrelated voxels. The denoised time series data was modelled using generalized linear modelling in SPM (GLM) after removing low frequency drift using a high-pass filter with cut-off of 128 s. In the main experiment, the activity of each voxel was modelled using 83 regressors (68 stimuli + 1 fixation + 6 motion regressors + 8 runs). In the localiser block, each voxel was modelled using 15 regressors (6 stimuli + 1 fixation + 6 motion regressors + 2 runs).

### ROI Definitions

All regions of interest (ROI) were defined using the data from functional localiser blocks together with anatomical considerations. Early visual areas (V1-V3) was defined as the region that responds more to scrambled objects compared to fixation. The regions identified were further parcelled into V1-V3 and V4 using anatomical masks from the *SPM Anatomy Toolbox* (Eickhoff *et al*., 2005). Lateral Occipital Cortex (LO) was defined as the voxels that responded to objects more than scrambled objects, but were restricted using the mask (Inferior Temporal Gyrus, Inferior Occipital Gyrus, and Middle Occipital Gyrus) created from tissue probability maps (TPM) labels available in SPM12. The Visual Word Form area (VWFA) was defined as a contiguous region in the fusiform gyrus that responds more to known words (Telugu/Malayalam) compared to scrambled words. The Temporal Gyrus (TG) was defined as voxels in the temporal gyrus (both superior and medial portions, and the Wernicke’s area) that responded more to known words (Telugu/Malayalam) compared to scrambled words. For each contrast, voxel-level threshold of p<0.001 (uncorrected) or cluster level threshold of p < 0.05 (FWE corrected) was used to define contiguous regions. However, for six subjects, VWFA could not be identified and therefore a lower threshold of p > 0.05 (uncorrected, lowest threshold p-value used was 0.2) was used until we observed a contiguous cluster of at least 40 voxels in Left fusiform gyrus. The LO and VWFA voxels were further restricted to the top-200 and top-20 significant voxels respectively (according to the T-value in the functional contrast). We obtained similar results with other choices of voxel selection. Finally, all results were visualized on the cortical surface using BSPMVIEW (http://www.bobspunt.com/bspmview/).

Example ROIs are shown in Fig. 4A for one subject, and a summary of their typical location and numbers of voxels is given in Section S7.

### Neural similarity in fMRI

For each ROI, the dissimilarity between each pair of stimuli was computed as 1 – r where *r* is the Spearman correlation coefficient between the activity patterns evoked across voxels by the two stimuli. The dissimilarities were z-scored and then averaged across subjects.

### Voxel population model

For each bigram, we modelled the response of a population of voxels as a linear combination of the response of the voxels to individual letters. For example, if there were 100 voxels in a given ROI, then for each bigram, its response is modelled as y = Xb, where y is 100×1 vector of beta (activation) values across voxels for that bigram, X is a 100×3 matrix, where first two columns correspond to the beta values for the corresponding voxels for the two constituent letters of the bigram, and the third column is a vector of 1s corresponding to a constant term, and b is a 3×1 vector of unknown weights that corresponds to the summation weights.

To evaluate model fit, we calculated the correlation between the observed and predicted response for each voxel. Doing so avoids the model fit from being biased by overall activation level differences between voxels. The correlation coefficients were averaged across all bigrams to obtain an average model correlation for that ROI in a given subject. The model fit was compared between readers and non-readers using paired t-test across subject-wise model correlations.

### Behavioral dissimilarity for fMRI bigrams

We estimated the behavioral dissimilarities for the bigrams used in the fMRI experiment using a reduced part-sum model. Recall that the part-sum model estimates separate letter dissimilarities for corresponding, across and within terms, but the estimated terms were all correlated with the single letter dissimilarities. We therefore reduced the part-sum model to a highly reduced model in which single letter dissimilarities from Experiment 1 combined linearly as follows.

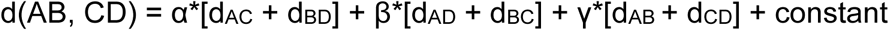

where d_AC_ d_BD_ etc are single letter dissimilarities observed from Experiment 1, and α, β, γ are unknown scaling terms for letter relations at corresponding locations, across-locations and within-bigrams. Thus this model has only four free parameters that can be estimated again using linear regression. To predict the behavioral dissimilarities between the bigrams used in the fMRI experiment, we first estimated the parameters of this reduced model from the bigram searches in Experiment 2, and then used these parameters together with the single letter dissimilarities from Experiment 1 to generate the predicted dissimilarities for all pairs of bigrams used in the fMRI experiment. This was then compared with the neural similarity calculated above.

### Experiment 4 (Bigram dissimilarity for fMRI bigrams)

To confirm the validity of this approach, we performed a separate experiment to measure perceived dissimilarity between bigrams used in the fMRI experiment. A total of 11 subjects (10 males, 24 ± 5 years, 7 Malayalam and 4 Telugu) participated in this experiment. Nine of these subjects participated in the fMRI experiment as well. Subjects performed searches for all pairs of 24 bigrams in each language, which amounted to a total of ^24^C_2_ x 2 = 552 unique searches. All other details were identical to Experiment 1. Subjects were highly consistent in their responses, as evidenced by a significant correlation between two halves of the subjects (Correlation in dissimilarity for odd- and even-numbered subjects: r = 0.51 & 0.66 for readers and non-readers of Telugu; r = 0.58 & 0.58 respectively for Malayalam; all correlations p < 0.00005). Critically, the observed dissimilarities were significantly correlated with the dissimilarities predicted from the reduced part-sum model and this correlation approached the consistency of the data itself (correlation for readers and non-readers: r = 0.65 & 0.75 for Telugu; r = 0.75 & 0.77 for Malayalam; all correlations p < 0.00005).

### Experiment 5 (Bigram fluency)

We performed an additional experiment to confirm the relation between reading fluency and bigram dissimilarity. A total of 22 subjects (7 female, 25 ± 6 years, 10 Malayalam and 12 Telugu) participated in this experiment. Subjects performed searches for all pairs of 25 bigrams of their native language, and also performed a reading fluency task. The 25 bigrams were the same as those used in Experiment 2. All other details were identical to Experiment 1. Subjects were highly consistent in their responses (Correlation between average dissimilarity of odd- and even-numbered subjects: r = 0.77 for Telugu readers on Telugu bigrams; r = 0.70 for Malayalam readers on Malayalam bigrams; p < 0.00005). The reduced part-sum model (described above) was used to estimate the strength of the corresponding, across and within terms. This model yielded excellent fits to the data, which approached or exceeded the split-half consistency of the data itself (model correlations: r = 0.84 for Telugu readers on Telugu bigrams, r = 0.85 for Malayalam readers on Malayalam bigrams, p < 0.00005).

## ACKNOWLEDGEMENTS

We are grateful to Mike Tarr, John Pyles and Elissa Aminoff for organizing an excellent fMRI workshop at IISc, and for help with standardizing scan and task parameters. These two critical efforts, which laid the groundwork for this study, were funded by a Tata Trusts grant and a CMU-IISc BrainHub grant (with SPA as co-PI).

## Author contributions

AA & SPA designed experiments; AA collected data and analysed data; AA, KVSH & SPA interpreted the data; AA & SPA wrote the manuscript with inputs from KVSH.

## Data and materials availability

All data necessary to understand and replicate the results are available in the manuscript or supplemental material. The behavioral and imaging data and the programs required to generate all the figures is available at https://github.com/iiscvisionlab/readingfMRI/

## Supplementary Material

### SECTION 1. STIMULI USED IN EXPERIMENTS 1-5

**Fig. S1.**
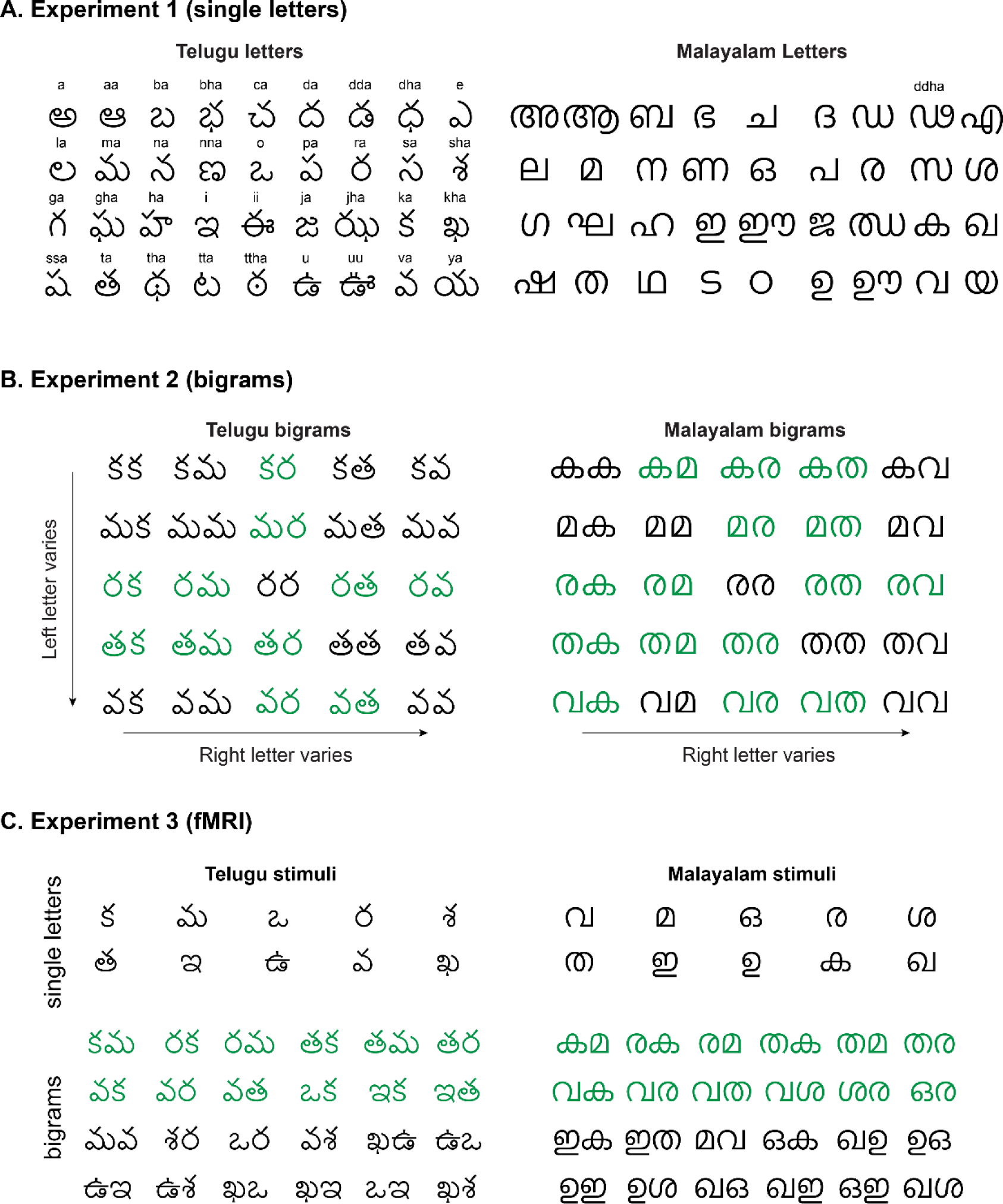
Stimuli used in Experiments 1-5. (A) Stimuli used in Experiment 1, together with their phonetic pronunciation which was identical in both languages despite the different letter shapes. One Malayalam letter was inadvertently different, but including/removing it made no difference to the results. Hence all results are reported including this one different letter. (B) Stimuli used in Experiment 2 & 5. Frequent bigrams in each language are highlighted in *green*. (C) Stimuli used in Experiment 3 & 4, comprising 10 single letters and 24 bigrams from each language. Frequent bigrams in each language are highlighted in *green*.

### SECTION 2. ADDITIONAL ANALYSIS FOR LETTERS (EXPERIMENT 1)

#### Visualization of perceptual space

In the main text, we found that pairwise letter dissimilarities are highly correlated between readers and non-readers, implying that reading only subtly alters letter representations. To examine whether these subtle alterations affect the overall structure of perceptual space, we visualized the letter dissimilarities using multidimensional scaling (MDS). Briefly, MDS finds the best-fitting 2D coordinates that best match with the observed distances. In the resulting plot, nearby stimuli correspond to hard searches.

The perceptual space obtained for Telugu letters is shown in Fig. S2A. Here, the MDS coordinates for letters in readers was shifted/rotated without altering their overall configuration so as to best match the MDS coordinates for letters in non-readers. Doing so enabled us to closely align the two representations. We observed no qualitative distortions between readers and non-readers (Fig. S2A). Likewise the perceptual space for Malayalam letters showed subtle differences but no overall distortion (Fig. S2B).

**Fig. S2.**
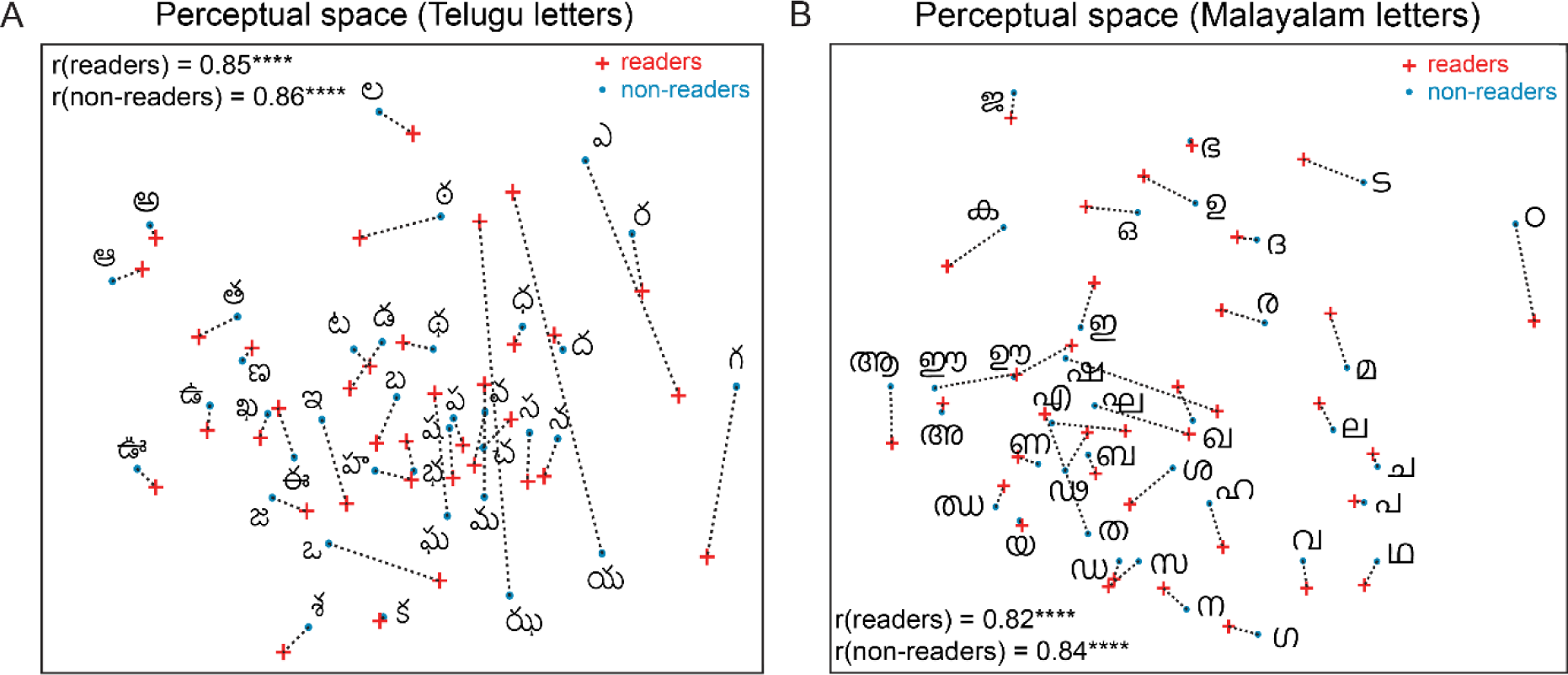
Reading expertise subtly alters perceptual space for single letters. (A) Perceptual space for Telugu letters. For readers, we used multidimensional scaling to find the 2D coordinates of all bigrams that best match the observed distances. In the resulting plot, nearby bigrams indicate hard searches. Red crosses correspond to readers and blue dots to non-readers. For non-readers, the multidimensional embedding obtained was shifted and rotated to match best with the embedding for readers (using the MATLAB function *procrustes*) for ease of comparison. Dotted lines indicate the change in each bigram with reading expertise. The correlation coefficient between dissimilarities in 2D plane and the observed data is shown for Malayalam and Telugu readers. Asterisks indicate significant correlation (**** is p < 0.00005). (B) Same as (A) but for Malayalam letters.

#### Bigram frequency analysis

To estimate bigram frequencies, we downloaded a text corpus of Telugu and Malayalam from HC Corpora (<u>corpora.heliohost.org</u>), which contains text from publicly accessible sources such as newspaper, twitter feeds, blogs etc. The text data was processed to remove unnecessary symbols/digits/English characters etc. Bigram frequencies from the text corpus was counted using the app “WordCreator” (www.sttmedia.com/wordcreator). Since Telugu and Malayalam languages have consonant modifiers (*maatras*), there were two choices for counting bigrams: to count occurrences of a pair of letters regardless of conjoined modifiers (e.g. రామ & రెమ are counted as an occurrence of రమ), or to count only occurrences of the exact pair without modifiers (e.g. only రమ is counted and not రామ or రెమ). In practice, however, both methods of counting yielded highly correlated bigram frequencies (r = 0.72, p < 0.00005 for Telugu and r = 0.79, p < 0.00005 for Malayalam) and therefore did not qualitatively alter the results. The bigram count of each bigram was converted into a fraction by dividing it with the summed count across all ^36^C_2_ = 630 letter pairs used in Experiment 1. This normalization is immaterial since it has no effect on the correlation coefficient.

To investigate the effect of bigram frequency, we asked whether the increase in dissimilarity in readers was related to the bigram frequency. This revealed a significant negative correlation across more frequent bigrams (r = −0.57, p < 0.005; Fig. S3A) and across all other Telugu searches (r = −0.12, p < 0.005). Thus, Telugu letters that cooccur together are perceived as more similar. This effect persisted even after accounting for the frequencies of single letters that comprised each bigram (partial correlations: r = −0.57, p < 0.05 across the 21 more frequent bigrams; r = −0.1, p < 0.05 across all other pairs). However, this effect was absent for Malayalam letters (r = 0.30, p > 0.05 for frequent bigrams; r = 0.03, p > 0.05 for all other pairs; Fig. S3B). This difference in correlation between Telugu and Malayalam was significantly different (p = 0.08 using a Fisher’s z-test comparing r = −0.12 with 609 pairs and r = 0.03 with 603 pairs; p = 0.002 comparing r = −0.57 with 21 pairs and r = 0.30 with 27 pairs). Thus, the effect of bigram frequency on visual search is not consistent across the two languages.

**Fig. S3.**
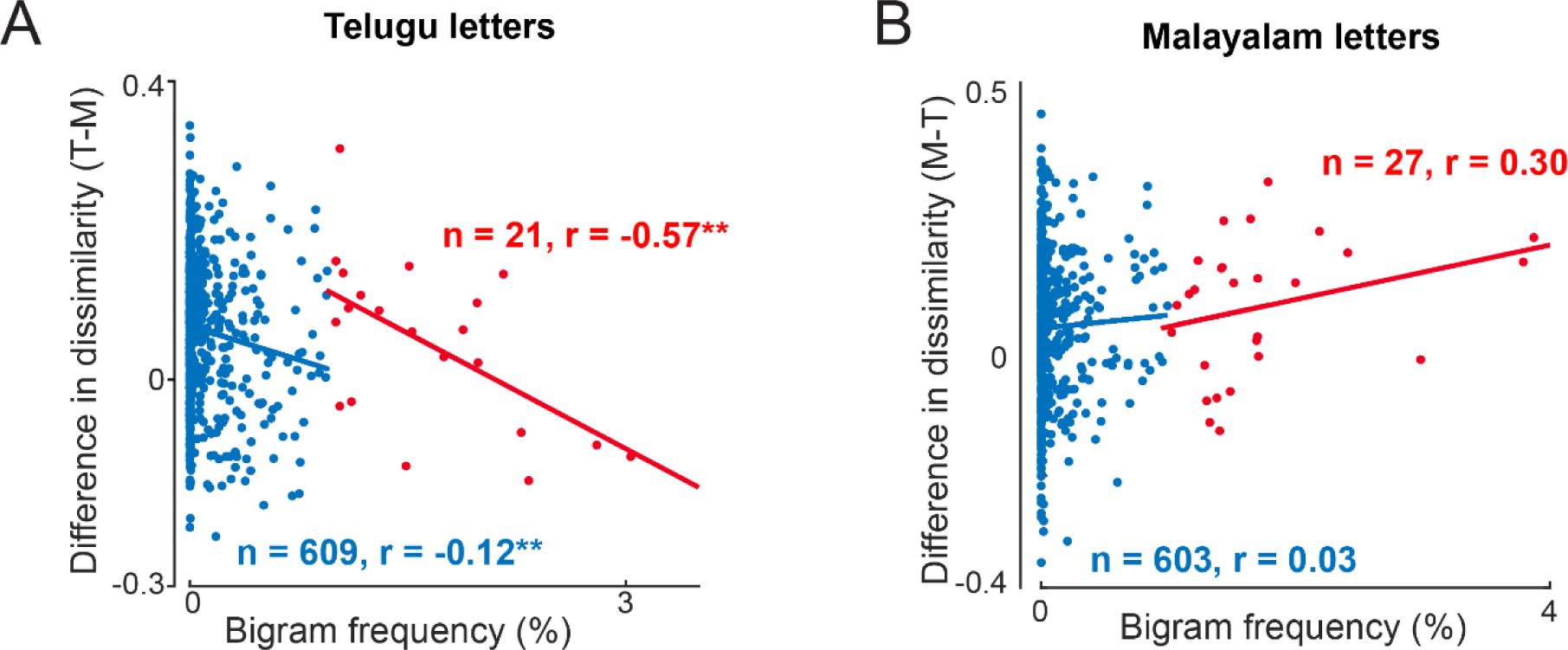
Effect of bigram frequency on perceived letter dissimilarities. (A) Difference in dissimilarity between readers and non-readers plotted against bigram frequency for Telugu letters. Blue points represent infrequent letter pairs (<1%) and red points represent frequent letter pair (>1%). Asterisks indicate significant correlation (**, p < .005). (B) Same as (A) but for Malayalam letters.

### SECTION 3. SOUND DISSIMILARITY (EXPERIMENT 6)

#### Introduction

The goal of this experiment was to characterize perceived sound dissimilarity for Telugu and Malayalam speakers. In particular we were interested in knowing whether sounds that occur frequently in bigrams tend to be perceived as more similar.

#### Methods

A total of 32 subjects participated in this experiment (19 male, 25 ± 4 years; 16 Telugu and 16 Malayalam). All 37 unique letters from the single letter experiment (Experiment 1) were selected and their pronunciation/sounds were downloaded from internet (http://web.cs.ucdavis.edu/~vemuri/classes/freshman/CourseOutline.htm). On each trial, two sounds were played one after the another through earphones, at the end of which subjects were asked to rate the dissimilarity between the two sounds by moving a sliding bar displayed on the screen, marked with “similar” and “dissimilar” at the left and right sides respectively. Subjects had the option to hear the same sound pairs multiple times before making the response and there were no time constraints.

#### Results

Both groups of subjects were highly consistent in their ratings, as evidenced by a high split-half correlation (r = 0.84 for Telugu, r = 0.88 for Malayalam, p < 0.00005 for both cases). Importantly they were equally correlated across groups as well (r = 0.91, p < 0.00005; Fig. S4A). To investigate the effect of bigram frequency, we asked if bigrams that were relatively more frequent in Telugu compared to Malayalam led to the corresponding two sounds being perceived as relatively less dissimilar in Telugu (or vice-versa). To this end we plotted the difference in sound dissimilarity rating (Telugu – Malayalam) against the difference in bigram frequency (Telugu – Malayalam). This revealed a significant negative correlation across the top 50 pairs with a large bigram frequency difference (r = −0.5, p < 0.0005; Fig. S4B). Thus, letters that co-occur in a given language tend to sound slightly more similar.

**Fig. S4.**
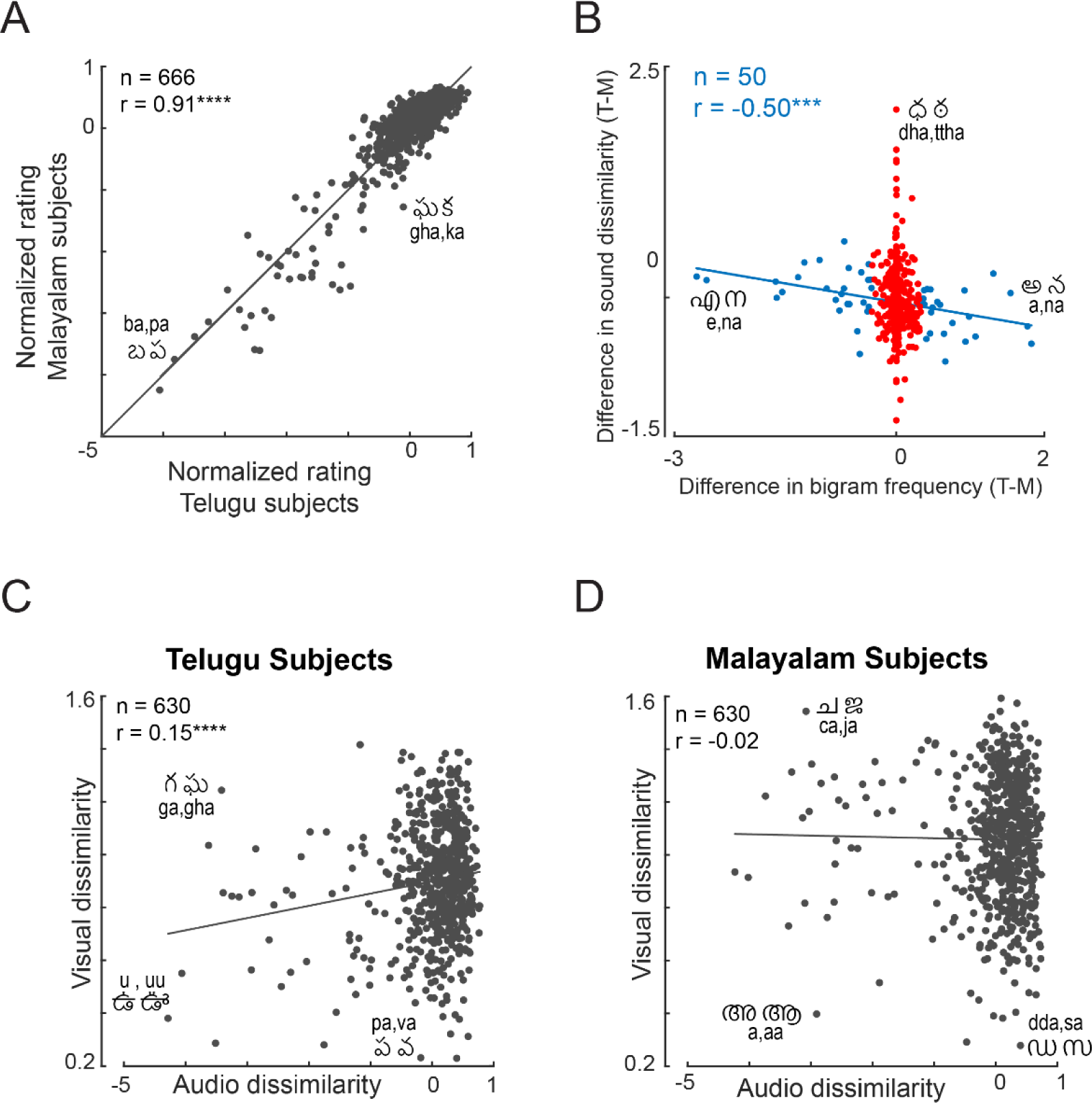
Effect of bigram frequency on perceived sound dissimilarity. (A) Correlation between perceived sound dissimilarity ratings between Malayalam and Telugu subjects. Asterisks represent statistical significance (**** is p < 0.00005). (B) Difference in z-scored sound dissimilarity rating (Telugu-Malayalam) plotted against the corresponding difference in bigram frequency for the two languages. Blue dots represent the top 50 pairs with large bigram frequency differences in the two languages. (C) Correlation between sound dissimilarity and visual dissimilarity for Telugu subjects, with example pairs highlighted. (D) Same as (B) but for Malayalam subjects.

Finally, although unlikely, we asked whether there was any correlation between sound dissimilarity and visual dissimilarity. A correlation might potentially exist since similar sounds may have been assigned by humans to similar shapes (for ease of remembering) or to dissimilar shapes (for ease of discrimination). However we found a weak correlation only for Telugu (Fig. S4C) but not Malayalam (Fig. S4D). This difference in correlation was statistically significant (p < 0.005, Fisher’s z-test).

#### Determining the best model for bigram search

In our analysis, we used inverse of reaction time (1/RT) instead of RT. This choice is motivated from our previous work which favoured 1/RT based models (Vighneshvel and Arun, 2013; Pramod and Arun, 2014). We verified it again in this study by fitting the part-sum model to either RT or 1/RT data and then testing the model prediction on RT or 1/RT data. We used two measures to evaluate model performance. The first was to compare the correlation coefficient between the observed and predicted data. The second was to compare models using a standard model selection criteria such as corrected Akaike information criterion (AICc). AICc for a model is calculated as AICc = N log (SS/N) + 2K + 2k(K+1)/(N-K-1). Here, N is the number of observations, SS is the sum of squared error between the model and data, and K is the number of free parameters in the model. A larger magnitude of AICc indicates a better model. The results for readers and non-readers on Telugu bigrams in shown in Table S1. It can be seen that 1/RT based models predict both RT and 1/RT data better compared to RT based models.

**Table S1.**
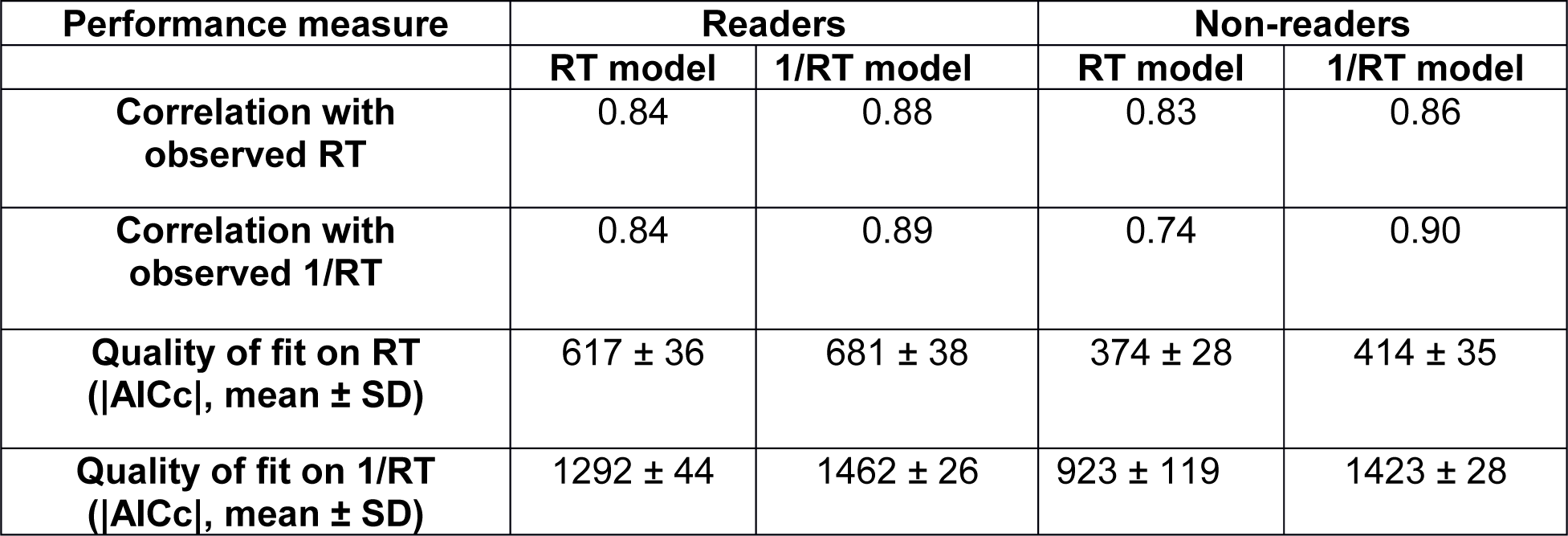
**RT vs 1/RT models.** We compared models fit to either observed RT or 1/RT, for their ability to predict RT or 1/RT. Larger AICc values indicate better fits.

Next, we compared the unique contribution of part relations at corresponding locations only, across locations only, and within locations only with the full part-sum model (Table S2). This revealed an interesting pattern: Corresponding-terms contribute more towards predicting dissimilarities for readers, while within-terms contribute more to the overall fit for non-readers. To quantify these contributions we performed a partial correlation analysis whereby we calculated the correlation between model predictions based on corresponding terms only after factoring out the model predictions based on the other two terms. This too revealed a similar pattern (partial correlations of each model term for readers after factoring out the other two: r = 0.69, 0.57 and 0.36 for corresponding, across and within-bigram terms, all correlations p < 0.00005). The better fit of the corresponding-terms-only model in readers reconfirms the finding that bigram representations are more compositional in readers.

**Table S2.**
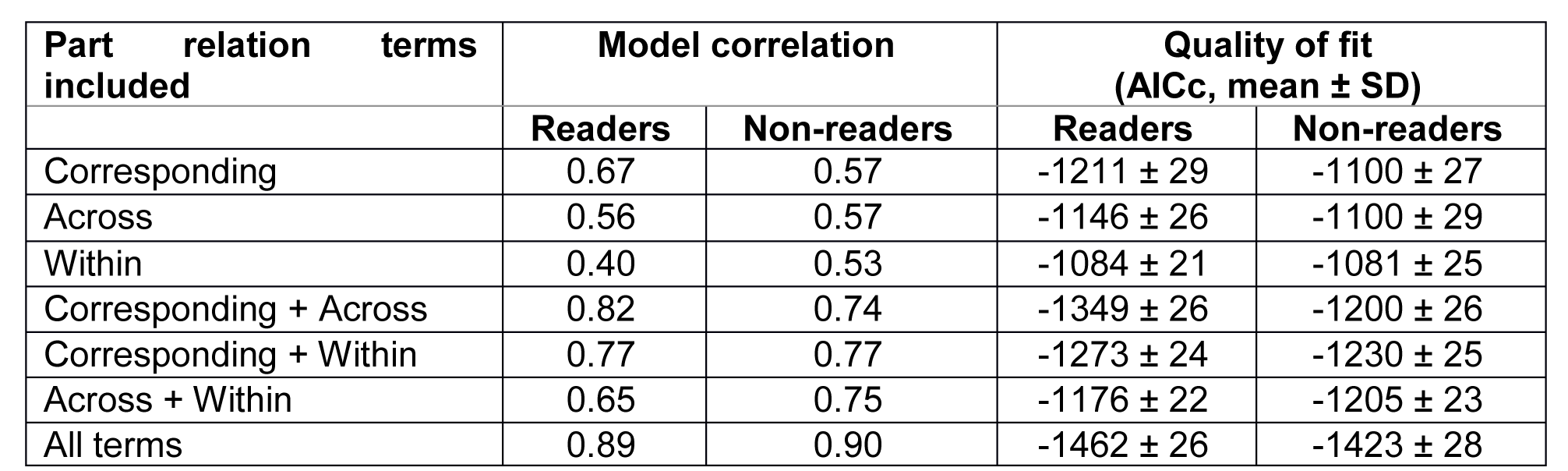
**Comparison of sub-models.** We compared both the overall correlation and quality of fit (absolute AICc) for part-sum models with only corresponding, only across or only within terms to compare their relative contributions.

| Part relation terms included | Model correlation | | Quality of fit (AICc, mean ± SD) | |
| --- | --- | --- | --- | --- |
|  | Readers | Non-readers | Readers | Non-readers |
| Corresponding | 0.67 | 0.57 | -1211 ± 29 | -1100 ± 27 |
| Across | 0.56 | 0.57 | -1146 ± 26 | -1100 ± 29 |
| Within | 0.40 | 0.53 | -1084 ± 21 | -1081 ± 25 |
| Corresponding + Across | 0.82 | 0.74 | -1349 ± 26 | -1200 ± 26 |
| Corresponding + Within | 0.77 | 0.77 | -1273 ± 24 | -1230 ± 25 |
| Across + Within | 0.65 | 0.75 | -1176 ± 22 | -1205 ± 23 |
| All terms | 0.89 | 0.90 | -1462 ± 26 | -1423 ± 28 |

#### First letter advantage in visual search

There is extensive evidence that there are letter position effects in recognizing letters in a word: the first letter is easier than the middle, and there is a characteristic W-shaped function along a word (Perea, Marcet and Gómez, 2016; Scaltritti, Dufau and Grainger, 2018). We therefore wondered whether there were similar effects in visual search. We first compared search times for bigrams that differed only in the left letter (e.g. AB vs CB, n = 50 pairs) with search times for bigrams differing in the right letter (e.g. BA vs BC, n = 50 pairs). To assess the statistical significance of these differences we performed an ANOVA on the search times with letter position (2 levels), search pair (50 levels) and subject (16 levels) as factors. For Telugu bigrams, both readers and nonreaders were faster to search between bigrams differing in the first letter compared to bigrams differing in the second letter (Fig. S6A). For Malayalam bigrams, this effect was present in readers but not nonreaders (Fig. S6B).

Next we asked whether this effect can be observed in the part sum model. However the part sum model described in the main text combined part differences at the left and right locations. We therefore fit a modified part sum model in which the net dissimilarity between a pair of bigrams is a sum of part relations at the left location, right location together with across-bigram and within-bigram terms as before. This modified part sum model contained 41 unknown parameters (10 each for the four types and a constant). The modified part sum model produced excellent fits of the data (correlation with observed dissimilarity for readers and nonreaders: r = 0.90 & 0.90 for Telugu bigrams, r = 0.92 & 0.93 for Malayalam bigrams; p < 0.00005 in all cases). Importantly, the left corresponding terms were significantly larger than the right side terms for readers of both languages (Fig. S6C,D). This effect was also present for Malayalam readers on Telugu bigrams (Fig. S6C) but not for Telugu readers on Malayalam bigrams (p = 1; Fig. S6D).

We conclude that reading expertise leads to a first-letter advantage in visual processing.

### SECTION 4. ADDITIONAL ANALYSIS FOR BIGRAMS (EXPERIMENT 2)

#### Visualization of changes in perceptual space

To visualize the effect of reading on bigram representations, we performed a multidimensional scaling analysis as in Fig. S2. The perceptual space for Telugu and Malayalam bigrams are shown in Fig. S5. It can be seen that the overall bigram representation is similar for readers and non-readers.

**Fig. S5.**
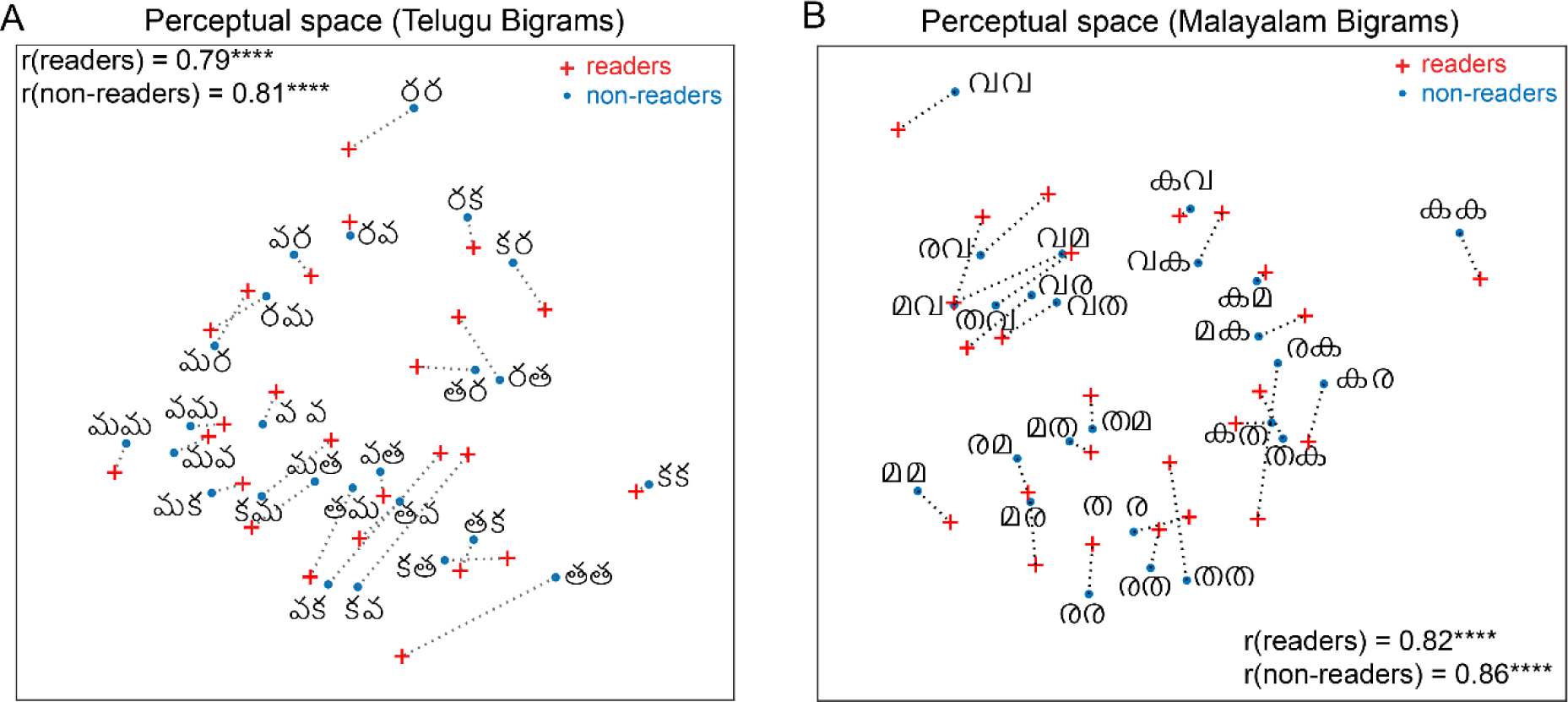
Perceptual space for bigrams in readers and non-readers. (A) Perceptual space for Telugu bigrams in readers and non-readers. For readers, we used multidimensional scaling to find the 2D coordinates of all bigrams that best match the observed distances. In the resulting plot, nearby bigrams indicate hard searches. Red crosses correspond to readers and blue dots to non-readers. For non-readers, the multidimensional embedding obtained was shifted and rotated to match best with the embedding for readers (using the MATLAB function *procrustes*) for ease of comparison. Dotted lines indicate the change in each bigram with reading expertise. The correlation coefficient between dissimilarities in 2D plane and the observed data is shown for Malayalam and Telugu readers. Asterisks indicate significant correlation (**** is p < 0.00005). (B) Same as (A) but for Malayalam bigrams.

**Fig. S6.**
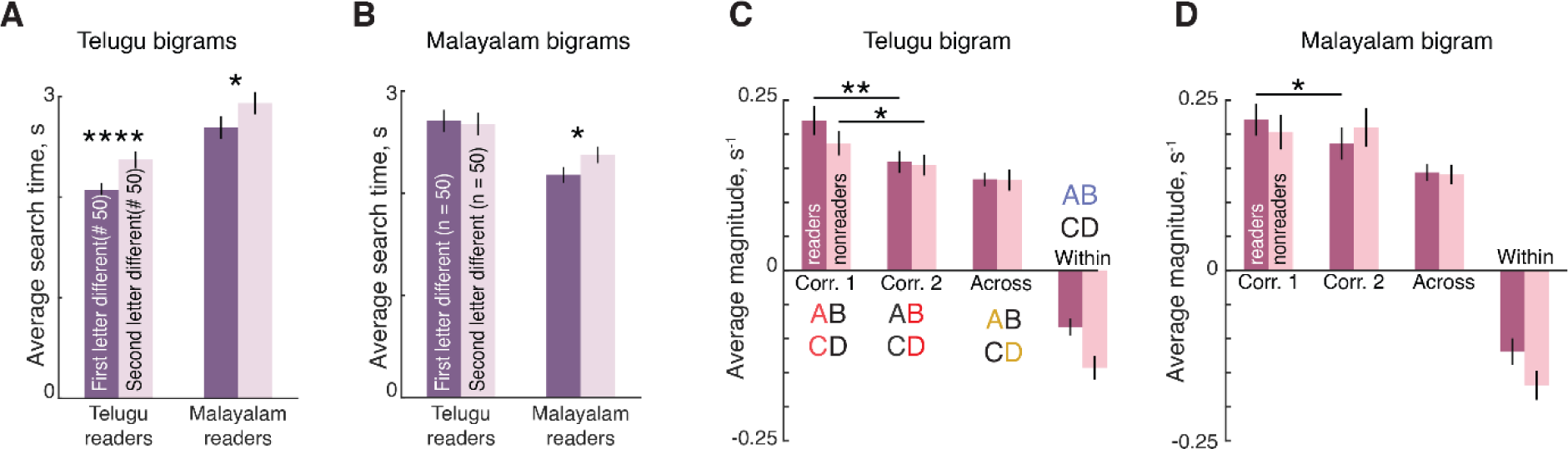
First-letter advantage in bigram search. (A) Average search times for search pairs where either the first letter was different (e.g. AB and CB, dark) or the second letter was different (e.g. AB and AC, light) for Telugu and Malayalam readers on Telugu bigrams. Error bars indicate standard error of the mean across pairs. Asterisks indicate statistical significance of the main effect of letter position in an ANOVA on RT with letter position, subject and search pair as factors (** is p < 0.005). (B) Same as (A) but for Malayalam bigrams. (C) Part sum model parameters (averaged across ^5^C_2_ = 10 part relations) for letter dissimilarities at corresponding location 1, corresponding location 2, opposite locations and within bigrams for readers (dark) and non-readers (light) for Telugu bigrams. Error bars indicate standard deviation across 10 model terms. Asterisks indicate statistical significance in a sign-rank test across 10 estimated part-part dissimilarities (* is p < 0.05, ** is p < 0.005, etc). (D) Same as (C) but for Malayalam bigrams.

### SECTION 5. TRIGRAM SEARCHES (EXPERIMENT 7)

#### Introduction

The goal of this experiment was to extend the findings of the bigram experiment (Experiment 2) to trigrams.

#### Methods

A total of 16 subjects (10 male, 23 ± 2 years; 8 Telugu and 8 Malayalam) participated in this experiment. Six single letters were chosen to create all possible (i.e. 6 x 6 x 6 = 216) trigrams each from Telugu and Malayalam. Unlike the bigram experiment, it was no longer feasible to sample all possible trigram pairs (^216^C_2_ = 23220). Instead, we sampled 325 pairs out of all possible search pairs such that each trigram appeared at least once and the resulting regression matrix of the part-sum model is not rank deficient. Of the 325 search pairs, there were 66 word-word pairs and 30 letter-transposition pairs. Thus subjects performed a total of 1300 correct trials (325 searches x 2 languages x 2 repetitions). All other details were identical to Experiment 1.

#### Results

An example trigram search array is shown using Telugu letters (Fig. S7A). As in the single letter and bigram experiments, we observed a clear double dissociation in search times (Fig. S7B) whereby subjects were faster for trigram searches in their known script.

**Fig. S7.**
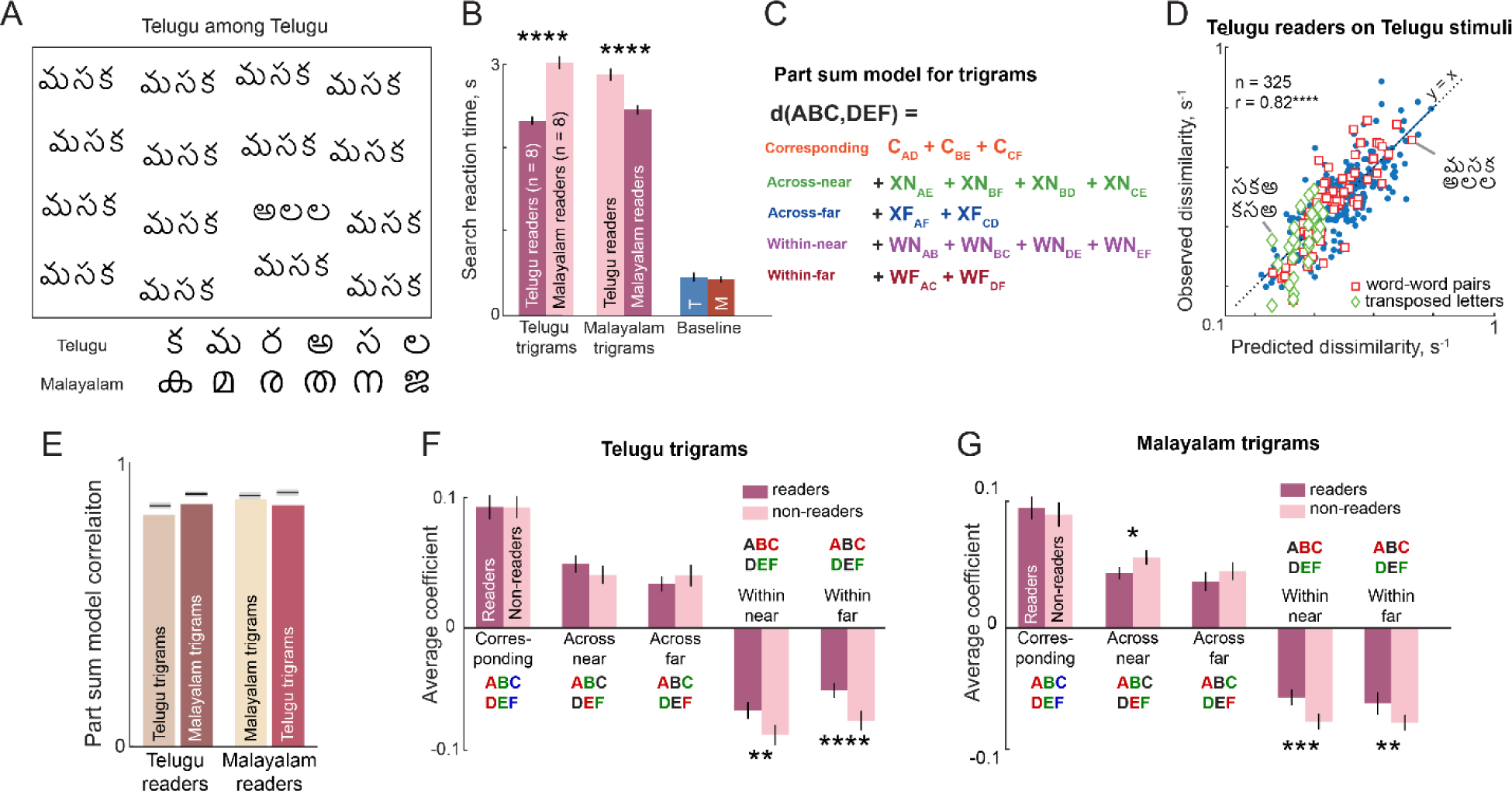
Part sum model for trigram dissimilarities. (A) Example trigram search array using Telugu letters. Below, Telugu and Malayalam letters (n = 6) used to create trigrams. (B) Average search time for readers and non-readers of Telugu letters & Malayalam letters, and response times for baseline trials. Asterisks indicate statistical significance (****, p < 0.00005). (C) Schematic of part summation model (see text). (D) Observed versus predicted dissimilarities for Telugu readers on Telugu bigrams. Each point represents one search pair (n = 325). Trigram pairs that comprised real words (*red diamonds*), trigram pairs with identical but transposed letters (*green circles*) are highlighted. Dotted lines represent the y = x line. Asterisks indicate statistical significance (****, p < 0.00005). (E) Part-sum model fit i.e. correlation between observed and predicted dissimilarities for readers and non-readers of Telugu and Malayalam. *Grey shaded bars* represent split-half correlation (after applying a Spearman-Brown correction) with 95% confidence interval. (F) Model parameters averaged across corresponding location terms, 15 near across location terms, 15 far-across location terms, 15 near-within trigram terms, and 15 far-within trigram terms for readers (*dark*) and non-readers (*light*) for Telugu trigrams. Error bars indicate standard deviation across 15 letter pairs. Asterisks indicate significant difference (****, p < 0.00005; ***, p < 0.0005; **, p < 0.005; *, p < 0.05). (G) Same plot as in (F) but for Malayalam trigrams.

We extended the part-sum model to predict the dissimilarities between trigrams. Specifically, the perceived dissimilarity between two trigrams ABC and DEF was modelled as a linear sum of all pairwise comparisons between letter pairs at corresponding locations (AD, BE, DF), letter pairs at near-across locations (AE, BF, BD, CE), letter pairs at far-across locations (AF, CD), nearby within-trigram pairs (AB, BC, DE, EF) and far within-trigram letter pairs (AC, DF). This is schematically shown in Fig. S7E.

Since all trigrams were created using 6 letters, there were a total of 76 parameters in the model (^6^C_2_ = 15 parameters each for corresponding, 15 for near across, 15 for far across, 15 for near within, 15 for far within part terms + 1 constant). The parameters were estimated by solving the linear equation **y** = **Xb**. Here, **y** is a 325×1 vector of observed dissimilarities, **X** is a 325×76 matrix whose rows contain either 0, 1 or 2 to indicate the absence, presence, or repetition of each term. **b** is the 76×1 vector that indicate the contribution of each part term. The vector **b** was estimated by solving linear regression.

#### Results

Subjects were highly consistent in their responses, as evidenced by a significant split-half correlation (correlation between odd- and even-numbered subjects for readers and non-readers: r = 0.63 & 0.71 for Telugu, r = 0.72 & 0.70 for Malayalam, p < 0.00005).

The part-sum model yielded excellent fits to the data, as illustrated for Telugu readers on Telugu trigrams (r = 0.82, p < 0.00005, Fig. S7D). As before we observed no systematic differences in residual error for frequent words or even transposed words(average residual error: 0.06 for frequent words, 0.06 for transposed words, 0.06 for others, p > .05 on a pair-wise rank sum test between the three groups). This correlation was nearly equal to the split-half correlation of the data, indicating that the model explained nearly all the explainable variance in the data. We obtained similar fits for readers and non-readers of both languages (Fig. S7E).

Once again, readers were faster than non-readers but their performance was explained entirely by the part-sum model. This meant that model parameters had to have changed to explain this difference. Indeed, we found as before that within-trigram interactions became weaker for readers compared to non-readers for both Telugu (Fig. S7F) and Malayalam (Fig. S7G) trigrams. We conclude that reading changes trigram representations by decreasing within-trigram letter interactions.

### SECTION 6. BIGRAM DISSIMILARITY & READING FLUENCY

To investigate the link between weaker letter interactions and reading fluency, we combined bigram dissimilarity and fluency data across Experiments 2, 4 and 5 (16 from Experiment 2, 11 from Experiment 4, 22 from Experiment 5). This yielded a set of 49 subjects (24 Telugu, 25 Malayalam). Data from a few subjects (n = 2) was excluded because they reported reading the passage multiple times to memorize it.

For each subject, we calculated the reciprocal of the average search time for all the bigram searches performed in that experiment. This observed dissimilarity was fit using the reduced part-sum model (see main text) to obtain the strength of the corresponding, across and within-bigram scaling terms as well as the constant term. Because each experiment contained different task contexts, we z-scored the model terms across subjects in each experiment before compiling them.

We calculated reading fluency from reading time as F = 500 – PRT, where PRT was the passage reading time in seconds. The number 500 was arbitrary but chosen to be larger than all passage reading times, and served simply to convert reading time (which is large when fluency is small) into a fluency measure (which is large when reading is fluent). Varying this number had no effect on the overall correlation. To remove language-specific fluency differences and experimental context differences, we z-scored the fluency data for each language and each experiment before combining it. The correlation remained robust across variations of these choices.

To investigate the link between reading fluency and bigram dissimilarity, we performed a regression analysis taking the corresponding, across and within-bigram terms as well as the constant term for each subject as regressors. This yielded a linear regression of the form **y** = **Xb** where **y** is a vector containing the reading fluency for each subject, **X** is a matrix with each term of the reduced part sum model along rows, and with number of rows equal to the number of subjects, and **b** is a vector of unknown weights reflecting the contribution of each term to the overall regression.

The observed and predicted fluency showed a significant correlation (r = 0.59, p < 0.00005; Fig. S8). The predicted fluency was influenced primarily by the within-bigram interactions and the constant terms, as evidenced by the fact that the model weights were roughly equal and that their confidence intervals did not include zero (model weights for corresponding, across, within and constant terms: 0.51, −0.12, 0.59 and 0.47 respectively; 95% confidence intervals for each: [-0.06 1.09], [-0.61 0.38], [0.19 0.99] and [0.14 0.80]). The contribution of within-bigram and constant terms were independent of each other, as evidenced by a significant partial correlation of each term while controlling for the other (Fig. 3H). While the within-bigram terms represent visual interactions between letters, the constant term reflects the overall motor speed of the subject. We conclude that reading fluency is influenced by both overall motor speed as well as visual processing.

**Fig. S8.**
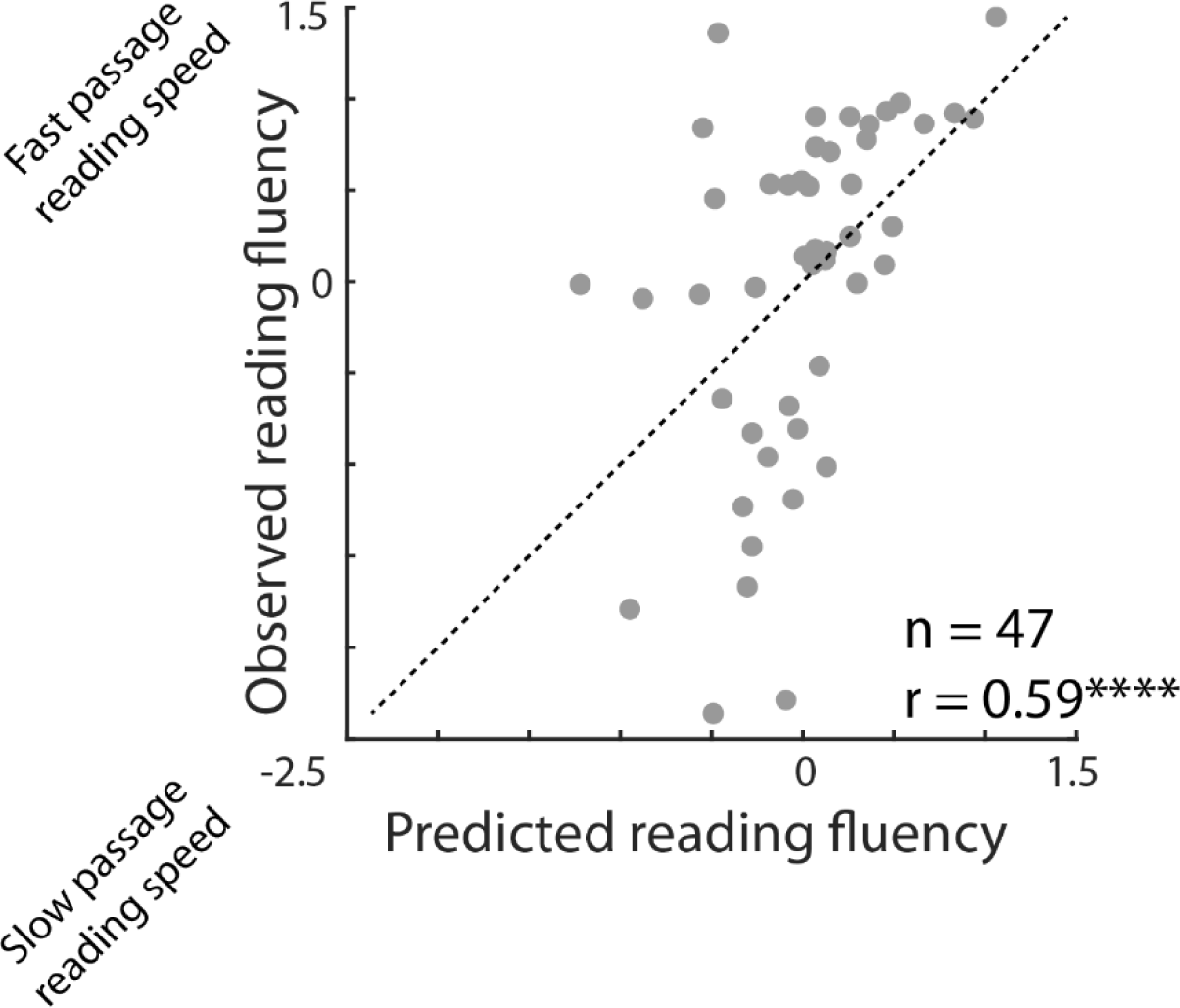
Within-bigram letter interactions correlate with reading fluency. Reading fluency in the fluency test for each subject plotted against the average within-term interaction terms estimated from bigram searches. Each point represents one subject. The correlation coefficient shown in the plot represents the Spearman’s rank-order correlation. The Pearson’s product-moment correlation for this data was likewise positive and significant (r = 0.49, p <0.0005).

### SECTION 7: ADDITIONAL ANALYSIS FOR EXPERIMENT 3 (FMRI)

#### ROI definitions

For each ROI, we report below the variability in its definition across subjects in terms of the number of voxels and the location of the peak in the normalized brain (Table S3).

**Table S3.** **Variability in ROI definitions across subjects.** For each ROI we report the mean and standard deviation across subjects of the number of voxels, and the XYZ location of the voxel with peak T-value in the normalized brain.

| ROI | Definition | #voxels<br>(mean ± sd) | ROI peak location |
| --- | --- | --- | --- |
| V1-V3 | Voxels activated for scrambled > fixation overlaid with anatomical mask of V1-V3 | 480 ± 171 | X: 1.7 ± 18.8<br>Y: -96.4 ± 4.4<br>Z: 5.7 ± 9.4 |
| V4 | Voxels activated for scrambled > fixation overlaid with anatomical mask of V4 | 228 ± 69 | X: 7.1 ± 22.8<br>Y: -87.3 ± 4.1<br>Z: 28.2 ± 11.5 |
| LO | Voxels activated for object > scrambled and not in other ROIs | 371 ± 125 | X: 6.8 ± 47.3<br>Y: -58.1 ± 11.1<br>Z: -21.1 ± 3.4 |
| VWFA | Voxels with known words > scrambled word in a contiguous region in fusiform gyrus | 81 ± 25 | X: -40.46 ± 13.9<br>Y: -48.6 ± 7.5<br>Z: -15.7 ± 4.9 |
| TG | Voxels with native words > scrambled word in a contiguous region in temporal gyrus | 388 ± 173 | X: -40.80 ± 41.2<br>Y: -45.1 ± 13.6<br>Z: 4.6 ± 6.4 |

#### Location of VWFA for Telugu and Malayalam

To compare the VWFA location across Telugu and Malayalam script, we calculated the number of subjects for which a particular voxel yielded a significant difference in the VWFA functional contrast within each group. The resulting activation maps are shown in Fig. S9. It can be seen that the location of the VWFA was similar for both Telugu and Malayalam readers.

**Fig. S9.**
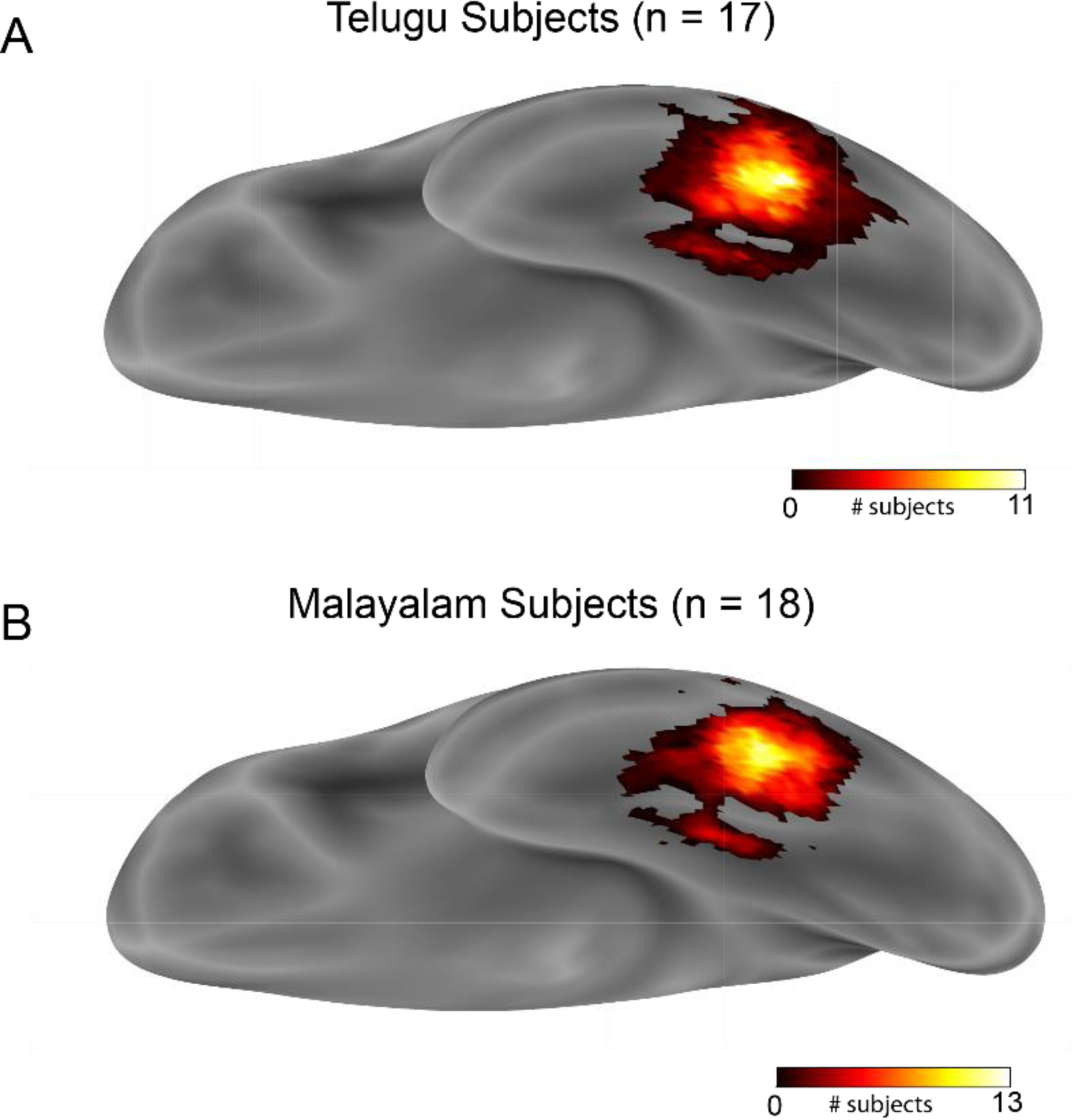
Location of VWFA. (A) VWFA in Telugu subjects. The color bar represents the number of subjects for which that particular voxel in the normalized brain showed difference in the words vs scrambled word functional contrast. (B) Same as (A) but for Malayalam subjects.

#### Neural dissimilarity in each ROI

For each pair of images, we calculated the neural dissimilarity in each ROI using the correlation distance measure, 1-r, where r is the Spearman’s correlation coefficient between voxel activations evoked by the two images for a given subject, and then averaged this dissimilarity across all subjects. The neural dissimilarity for all pairs of images in each ROI is depicted in Fig. S10.

**Fig. S10.**
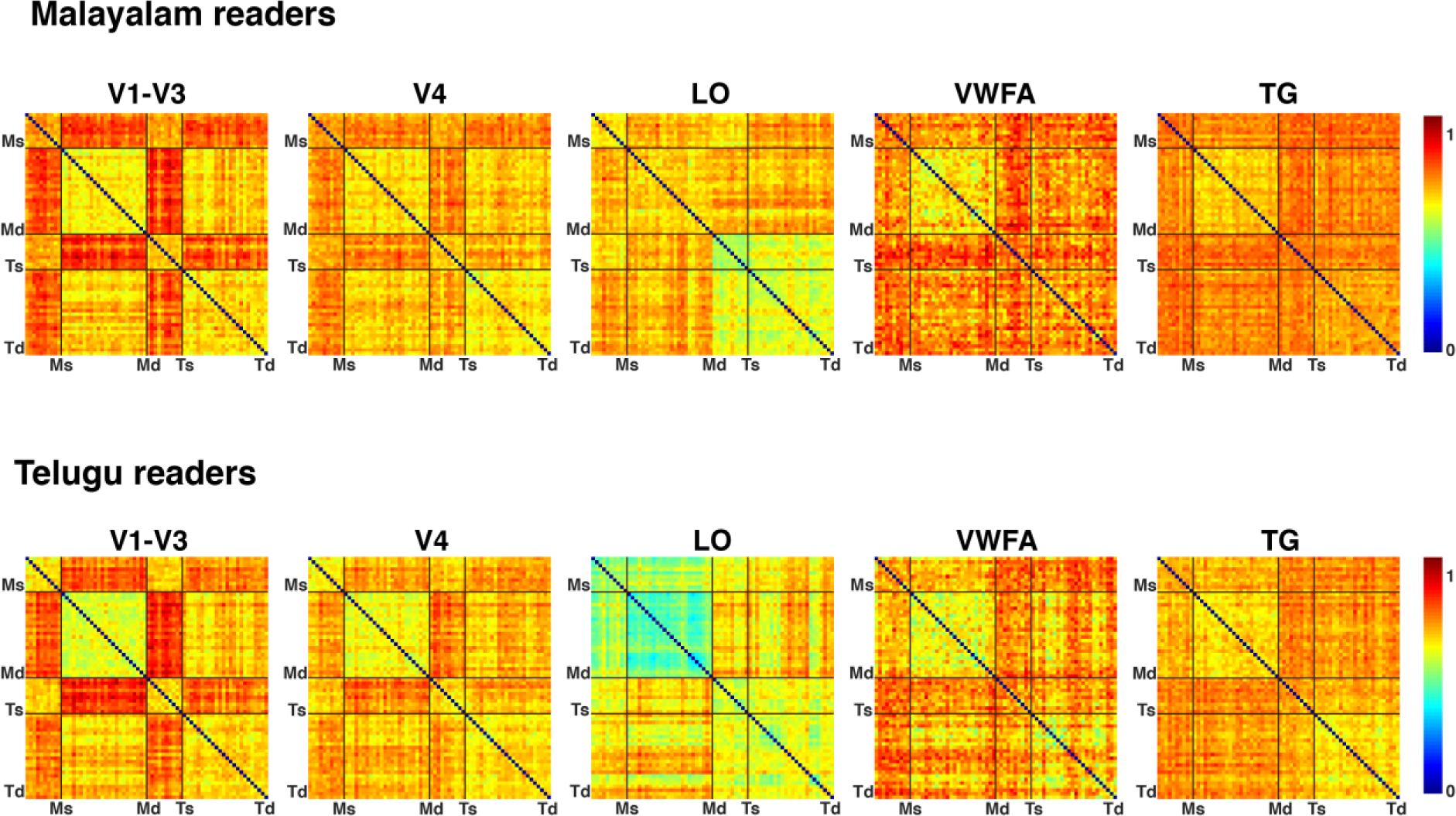
Representational dissimilarity matrix across ROIs. (A) Averaged pair wise dissimilarity across Malayalam readers for all possible pairs of stimuli. Same analysis is performed using voxels from V1-V3, V4, LO, VWFA, and TG. Colour bar indicates the dissimilarity value i.e. 1 – r (Spearman correlation coefficient across activity pattern in a given ROI). Ms – end of Malayalam single letter stimuli, Md - end of Malayalam double letter stimuli, Ts – end of Telugu single letter stimuli, Td – end of Telugu double letter stimuli. (B) Same plots as in (A), but for Telugu readers.

#### Searchlight analyses

To identify other brain regions that might show the effects observed in the individual ROIs, we performed a whole-brain searchlight analysis. Specifically, for each voxel in a given subjects’ brain, we considered a local neighbourhood of 27 voxels (3×3×3 voxels) and performed the following analyses of interest. We obtained similar results for larger searchlight volumes.

##### Searchlight for activation differences for known versus unknown scripts

For each voxel, the betas for the known stimuli (median across all the known stimuli and followed by median across all subjects) was subtracted from the median betas for the unknown stimuli and divided it with the standard deviation of the difference to estimate T value. Overall, Lateral Occipital Cortex (LO), Superior Parietal Lobe (SPL), and pre-central gyrus (PCG) responded more to the unknown stimuli, while language areas and visual areas along the ventral surface along the fusiform gyrus responded more to the known stimuli (Fig. S11).

**Fig. S11.**
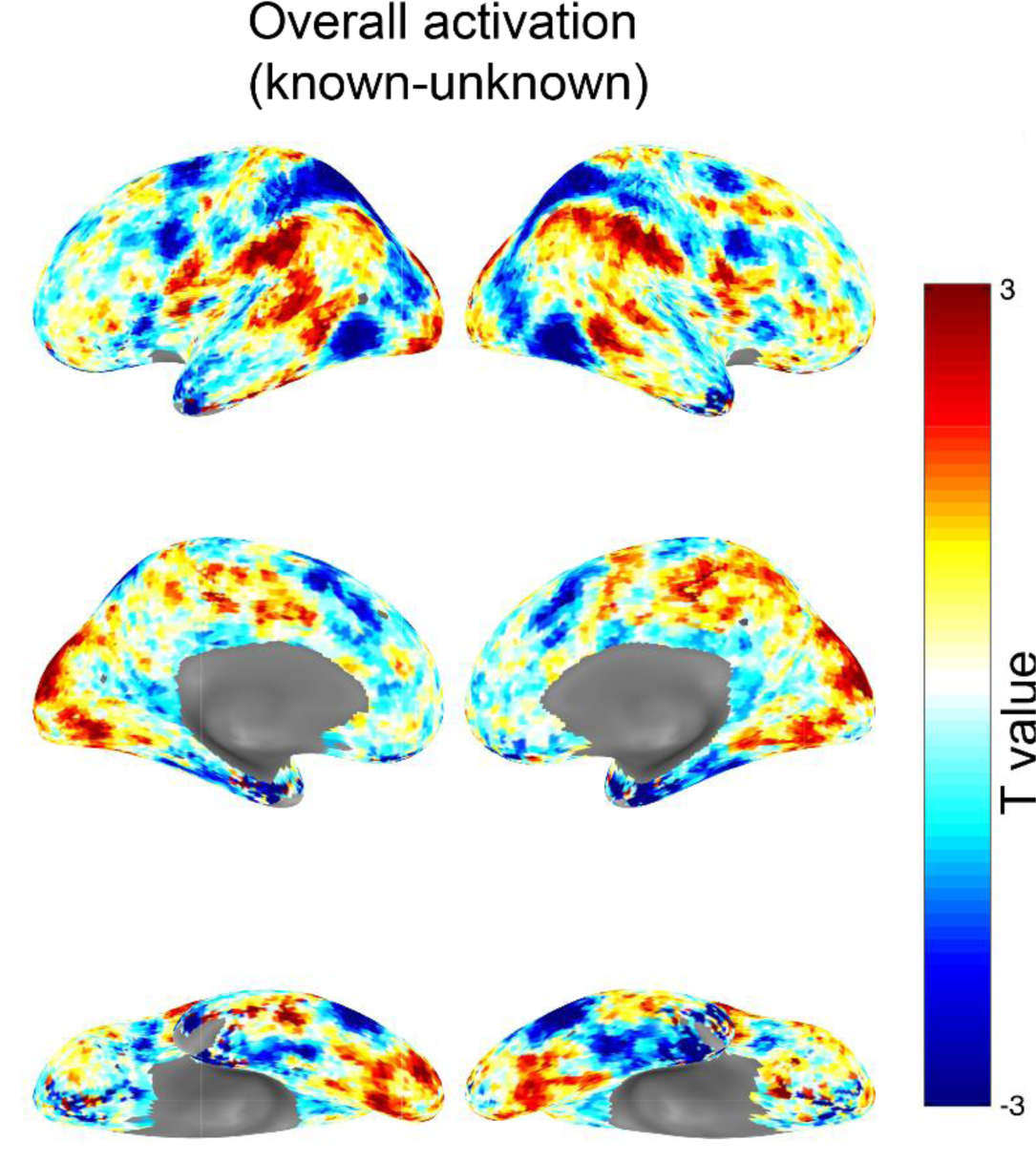
Differences between known and unknown scripts. Searchlight map of depicting difference between known and unknown script averaged across subjects. *Searchlight for regions where neural dissimilarity matches with behaviour*. For the neighbourhood of each voxel, we calculated the pairwise neural dissimilarity for all bigrams for a given subject, and averaged this across subjects. We then calculated the correlation between this local neural dissimilarity and the corresponding bigram dissimilarities predicted from the model, separately for bigrams of known and unknown scripts. This correlation map was visualized on the brain surface for known (Fig. S12A) and unknown (Fig. S12B) scripts. It can be seen that behavioural dissimilarities for known scripts match best with a region centred around LO as well as in temporal gyrus and parietal/motor regions (Fig. S12A). By contrast, for unknown scripts the peak activation is more posterior, closer to the early visual areas (Fig. S12B).

**Fig. S12.**
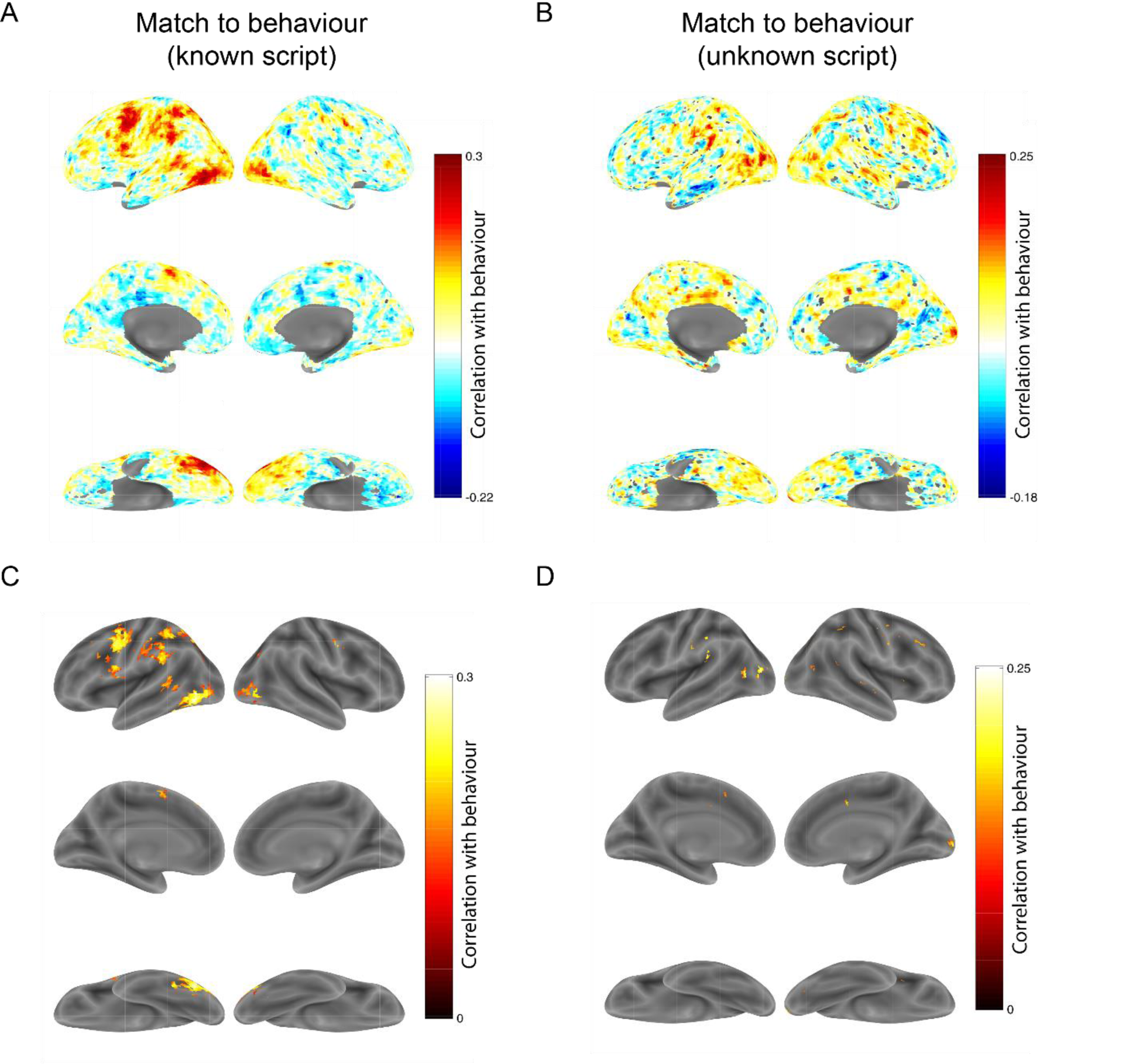
Neural activations that match with behaviour for known and unknown scripts. (A) Searchlight map of correlation between neural dissimilarity and bigram dissimilarities in behaviour for known scripts. (B) Same as (A) but for unknown scripts. (C) Same as (A) but with only statistically significant correlations (p < 0.05). (D) Same as (B) but with only statistically significant correlations (p < 0.05).

##### Comparing part-summation model fit for the known and the unknown stimuli

For each subject and each local voxel neighbourhood, we modelled the responses of each bigram across voxels as a linear combination of the responses of its constituent letters. The model fit (correlation between observed and predicted betas across bigrams) was evaluated for the central voxel in the mask for known and unknown scripts separately. We then calculated the difference in model correlation for known – unknown scripts and smoothed this map using a Gaussian with FWHM of 5 mm. The resulting searchlight map shows that, within the visual regions, bigram responses were more predictable from single letters for known compared to unknown scripts in the anterior and ventral portion of the LO as well in the fusiform gyrus and parietal cortex (Fig. S13A). One subject with very low reading fluency was excluded from this analysis.

**Fig. S13.**
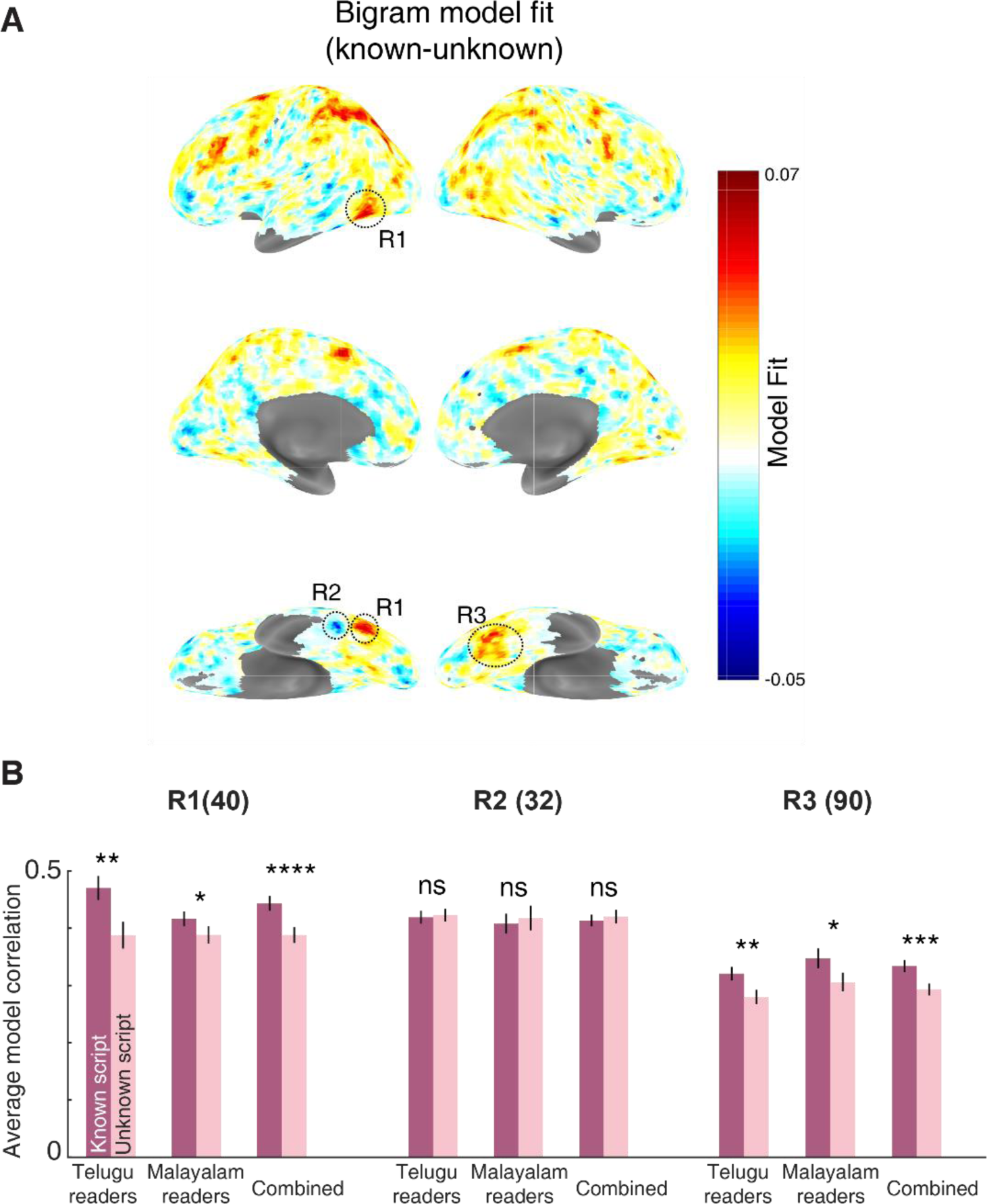
Difference in model fit for known versus unknown scripts. (A) Searchlight map depicting the difference in model fit for known versus unknown scripts for each voxel, averaged across subjects. Three regions in the visual cortex that showed differences between known and unknown scripts are highlighted: region R1 is located on the anterior-ventral portion of left LO, region R2 is located in the left fusiform gyrus close to the VWFA and R3 is located in the right fusiform gyrus. (B) Average model correlations for known and unknown scripts using split-half analysis (see text) for all three regions (R1, R2, R3) for Telugu readers, Malayalam readers and both combined. Numbers in parentheses represent the average number of voxels in each region. Asterisks indicate statistical significance obtained by comparing subject-wise average model fits using a sign-rank test (* is p < 0.05, ** is p < 0.005, and so on).

To assess the significance of these effects, we performed an ROI analysis on the above three regions in the visual cortex, marked as R1, R2, and R3 in Fig. S13A (normalized peak coordinates: R1: -39,-61,-10; R2: -45,-43,-19; R3: 36,-58,-13). If we simply compared model fits in these regions directly it would be circular since these regions were identified by the same criterion. To avoid this, for each region e.g. R1, we repeated the searchlight analysis on odd-numbered subjects, identified the region R1, and then selected these same voxels from even-numbered subjects to compare model fits, and vice-versa. We averaged the results from R1 voxels from even-numbered subjects and R1 voxels from odd-numbered subjects. We did likewise for R2 and R3.

This split-half analysis ensures that, if these regions were simply noise, there would be no net difference in model performance on average. However this was not the case: model performance was indeed better on known scripts compared to unknown scripts in region R1 and R3 but not in region R2 (Fig. S13B).

Thus, reading expertise leads to more compositional bigram representations in the anterior-ventral portion of the lateral occipital cortex and fusiform gyrus.

#### Effect of bigram frequency

Although previous studies have shown frequency related effects in VWFA (Kronbichler *et al*., 2004), we did not find any study that directly compared the activity of frequent and infrequent bigrams in VWFA. Here, we equally grouped the 24 bigrams into high frequency (HF) and low frequency (LF) bigrams. For each language, we compared the mean activity of HF and LF bigrams for both readers and non-readers (Fig S14A,C). Non-readers served as a baseline measure and accounted for any stimulus dependent effects. We observed a significant difference in mean activation levels between low and high frequency bigrams only for Telugu letters in LO & VWFA. However these differences were abolished upon factoring out single letter frequencies (Fig. S14B,D). Thus, bigram frequency does not modulate activity in LO or VWFA over and above that expected from single letter frequency. These results also demonstrate that it is critical to control for individual letter frequencies before invoking any bigram effects.

**Fig. S14.**
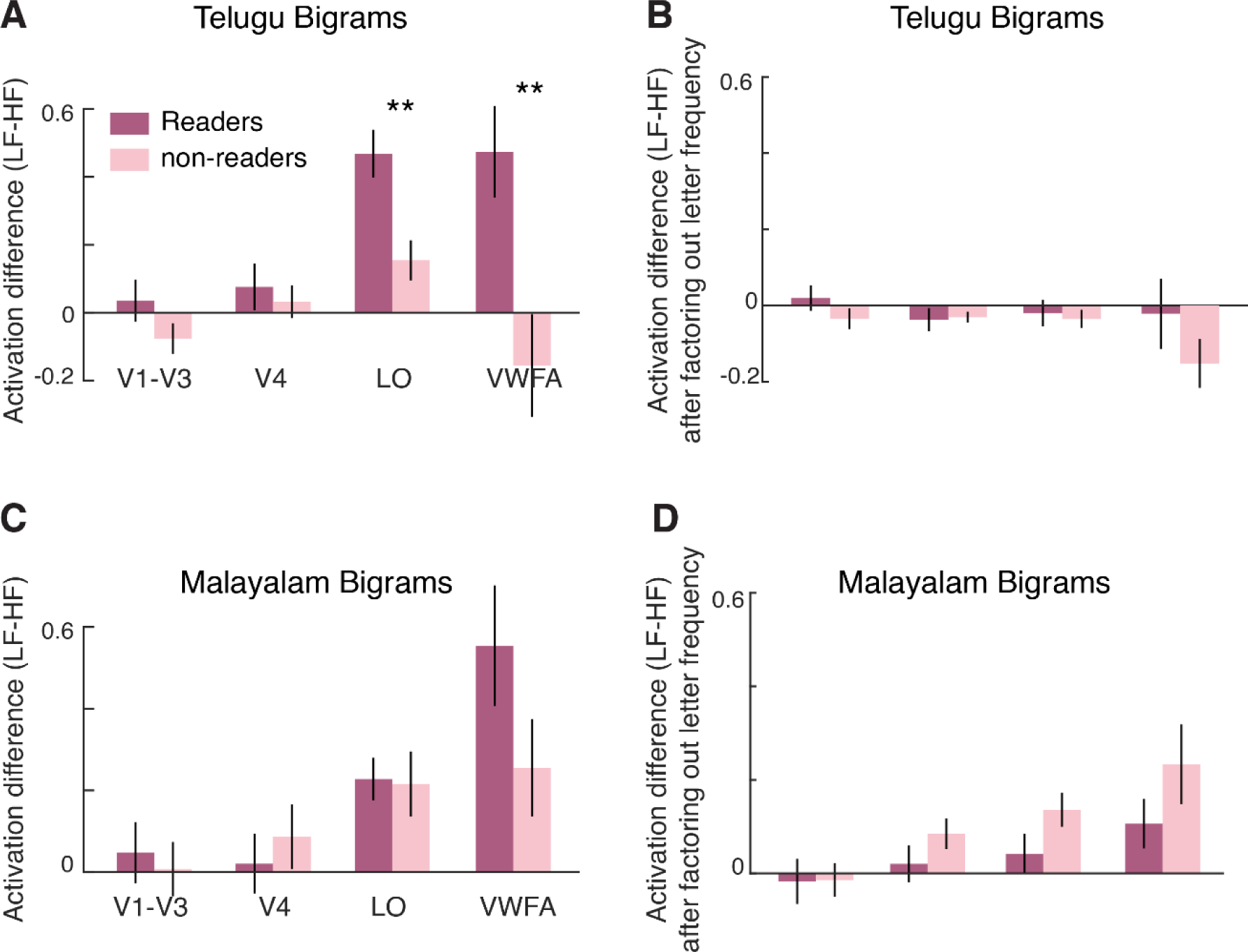
Effect of bigram frequency in each ROI. (A) Average difference in activation between LF and HF bigrams in each ROI. Asterisks indicate statistical significance calculated on mean activations across each subject (** is p < 0.005). (B) Same as (A) but after factoring out the effect of letter frequency. To do so, we fit a linear model where the mean activation of each ROI to a given bigram is a linear sum of the first letter frequency, second letter frequency and a constant term. The predicted activation of this letter frequency model was subtracted from the observed activation, and then we compared the residual unexplained activation for low-frequency and high-frequency bigrams. In the resulting plot, no comparison was statistically significant. (C) Same as (A) but for Malayalam bigrams. (D) Same as (B) but for Malayalam bigrams.

**Fig. S15.**
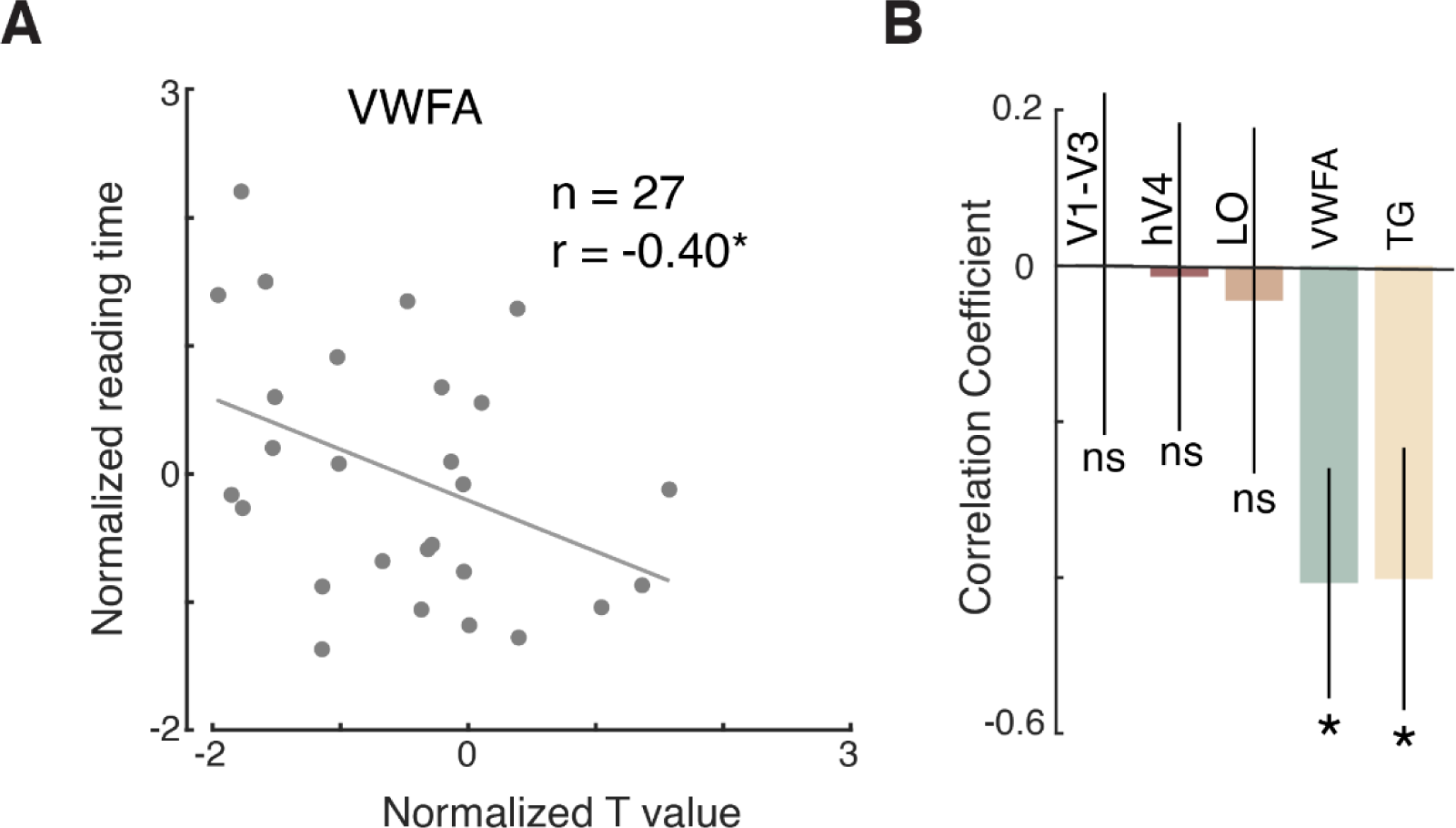
VWFA and reading fluency. (A) Correlation between normalized reading time and average normalized T values across voxels of VWFA. Each point represents one subject. Asterisks indicate significant correlation (* is p < .05). (B) Correlation between normalized reading time and normalized T value (averaged across voxels in a given ROI). Error bars represent standard deviation of this correlation calculated across 100 bootstrap samples where subjects were repeatedly chosen randomly with replacement. Asterisks represent the fraction *p* of bootstrap samples in which the correlation was smaller than zero (* is p < 0.05).

The above results show that bigram frequency effects do not modulate overall ROI activations but they could still influence smaller subsets of voxels in the ROI. To assess this possibility we additionally performed a voxel-wise analysis. For each voxel, we calculated the partial correlation between its mean activity and bigram frequency after regressing out the contribution of the two single letter frequencies in each bigram. This revealed 3.7% of the voxels (across all subjects) in LO and 3.9% of all VWFA voxels with a significant partial correlation (p < 0.05). Since these fractions of voxels are below the level expected by chance, we conclude that bigram frequency does not modulate LO or VWFA activation even at the single voxel level.

Finally, we note that some studies have reported increasing response of VWFA to word-like stimuli (Binder *et al*., 2006; Vinckier *et al*., 2007) whereas others have reported the exact opposite trend (Kronbichler *et al*., 2004). We speculate that these disparate effects can be reconciled if one accounts for single letter frequency and position effects.

#### Correlation between VWFA and reading fluency

In attempt to validate our results with previous observations, we correlated normalized reading time (z-scored across subject within each group) with the normalized T-value (estimated from the localiser block) averaged across voxels from VWFA. As reported in previous studies (Dehaene *et al*., 2010), we found negative correlation between reading time and VWFA activity. We excluded 8 subjects from the analysis (2 subjects did not participate in the fluency test, 4 subjects did not follow the instructions of reading task and read the passage multiple times before responding, and we could not localise VWFA for 2 subjects). To ensure that the results were not driven by few subjects, we repeated the above analysis across 100 bootstrap samples. Only VWFA and TG regions showed significant correlation between reading time and activity.

